# Development, validation, and evidence of frequent exchange of mating type *HD* alleles of *Puccinia striiformis* f. sp. *tritici* provide an insight into highly genetic diversity

**DOI:** 10.1101/2025.05.06.652357

**Authors:** Sinan Li, Mudi Sun, Zhiming Du, Xiangrui Cheng, Letao Yang, Meng Ju, Zhensheng Kang, Jie Zhao

## Abstract

Wheat stripe rust, caused by *Puccinia striiformis* f. sp. *tritici* (*Pst*), is a worldwide destructive wheat disease. It possesses the potential to generate novel virulence races through sexual reproduction, which is the primary cause of its extensive epidemics. Mating type (MAT) of *Pst* lacks direct experimental evidence to account for the role of sexual recombination on high genetic diversity in evolution. Herein, we developed 11 pairs of *HD* alleles of *Pst* MAT genes, and 2 pairs *HD* alleles of *P. striiformis* f. sp. *hordei* (*Psh*, the causal agent of barley stripe rust) MAT genes. *HD* alleles in *Pst* are different from those in *Psh*. We tested that mating system of *Pst* is tetrapolar responsible for sexual genetic recombination via *HD* genes by experiments of selfing and sexual hybridization on alternate (secondary) hosts. Multiple *HD* gene types and *HD* gene recombination types were detected in Chinese natural *Pst* populations of 200 *Pst* isolates and 13 *Psh* isolates from 92 sampling sites in 11 provinces, supporting that sexual reproduction accounts for frequent emergence of new races and high genetic diversity of *Pst* population in northwestern oversummering variable region (hotspot) than other stripe rust-occurring regions in China. Overall, these results provide an insight into diverse types and frequent exchange of *Ps* MAT alleles, revealing the important role of sexual recombination via MAT alleles in virulence variation, high genetic diversity and epidemics of *Pst*, and also provide a basis for wheat stripe rust control in those regions where sexual reproduction occurs.

**Author summary:** Heteroecism of the wheat stripe rust pathogen *Puccinia striiformis* f. sp. *tritici* (*Pst*) was discovered a decade ago. The significance of sexual reproduction in maintaining genetic diversity has emerged as a pressing concern in contemporary scientific discourse. In this study, we elucidated the presence of multiple *HD* gene types (*HD1* linked to *HD2*) in *Puccinia striiformis* f. sp. *tritici* (*Pst*), distinguishing it from *P. striiformis* f. sp. *hordei.* Through biological experiments on sexual reproduction, we demonstrated that the mating system of *Pst* is tetrapolar. Furthermore, we observed that sexual reproduction, particularly selfing (individual race) and crossing (between two races), facilitates sexual recombination via *HD* genes, leading to pathogenic variation and a high level of genetic diversity. We established a model illustrating sexual reproduction of *Pst* by selfing and crossing, respectively. In China, we have observed that the *Pst* population exhibits a diverse range of *HD* gene types and *HD* gene recombination types. Notably, northwestern oversummering variable regions (hotspot) have a higher number of *HD* genes compared to other epidemic region. Population structure and cluster analyses based on *HD* genes indicate that *Pst* populations in Tibet and Xinjiang are distinct from those in other inland regions. These findings provide valuable insights into comprehending the intricate interplay between sexual reproduction and genetic diversity of *Pst*, thereby elucidating the reasons behind the high genetic diversity of the Chinese *Pst* population.

## INTRODUCTION

Wheat (*Triticum aestivum*) is one of staple food crop worldwide. It is vulnerable to wheat stripe (yellow) rust, caused by *Puccinia striiformis* f. sp. *tritici* (*Pst*). The disease poses a destructive threat, capable of causing substantial yield reductions during severe epidemic years. In the event of an early and severe infection affecting highly susceptible wheat cultivars under favorable conditions, it is possible to experience complete crop failure [1–5]. In China, since the 1940s, eight severe outbreaks of wheat stripe rust have been documented. The most severe epidemic occurred in 1950, affecting 10 million hectares of wheat and resulting in approximately 6 million metric tons of wheat loss. [6–9].

In agricultural practices, cultivating genetically resistant wheat varieties serves as a fundamental mechanism for controlling wheat stripe rust. However, new races frequently emerge that overcome the resistance of wheat cultivars, rendering them susceptible within a span of one to ten years after their release. Some *Pst* races develop rapidly and become predominant, resulting in subsequent large-scale epidemics that often can cause remarkable yield reductions across the world [2,4,10–12]. Historically, due to emergence of new races and rapid accumulation, eight replacements of mainly cultivated wheat varieties nationwide have reported in China [7,12–13]. Similarly, in Europe new races such as ‘Warrior’, ‘Kranich’, and ‘Triticale aggressive’, possibly originated from sexual recombination with non-European *Pst* population, colonized and replaced local stripe rust populations after 2011 due to their rapid spread [14].

In contrast, the Chinese *Pst* population exhibits a high level of genetic diversity, whereas the rust populations in America, Europe, and Australia are characterized as a clonal [10, 15–21]. For example, the genetic diversity of a *Pst* population in Tianshui City, Gansu Province, was reported within a single year to be over one thousand times greater than that of French populations over a twenty-year period [18]. In the context of distinguished population genetic diversity, genetic recombination of Chinese natural *Pst* population through sexual cycle was strongly speculated to exist in the context of its sexual stage that was unknown [17–18]. Subsequently, sexual stage of the rust, which occurs on barberry (*Berberis*), was discovered in 2010 [22], and this speculation was proved by many *Berberis* species growing in different regions of China in multiple independent years [23–27]. More importantly, susceptible barberry can serve as source of aeciospore inoculum of *Pst* to nearby wheat fields and causing stripe rust, playing a role in occurrence of the disease in those areas where susceptible barberry and wheat coexist under natural conditions in China [25–27]. Consequently, it is responsible for highly genetic diversity of Chinese *Pst* population.

Sexual reproduction is a fundamental process for the genetic recombination of fungi and the biological evolution of species. It is one of the defining characteristics of biological diversity in nature [28–29]. It can produce diverse genotypes, facilitating the capacity of fungal species to environmental adaption and selection [30–31]. Fungal sexual reproduction is mainly controlled by mating type (MAT) genes responsible for regulating gamete and hormone of bisexual affinity cells [32–34]. MAT genes play a vital role in genetic evolution of fungi [34]. Based on a MAT gene in a haploid cell, fungi complete sexual reproduction by means of heterothallism, homothallism, or pseudo-homothallism. Based on the number of pairs of incompatible factors controlling MAT, heterothallism is classified into bipolar and tetrapolar heterothallism. A fungus of bipolar heterothallism has two types of MAT genes which is controlled by a pair of allele loci, and that of tetrapolar heterothallism has four types of MAT genes being controlled by two pairs of allele loci [35–37], viz. *A* MAT gene locus and *B* MAT gene locus [30, 36, 38]. *A* MAT gene locus encodes two types of homeodomain transcription factors (*HD* gene locus), and *B* MAT gene locus encodes lipopeptide pheromones and pheromone receptors (*P/R* gene locus). *A* MAT gene locus includes two types of alleles that encode homeodomain transcription factors, *HD1* and *HD2* genes [39–40]. The product of *HD1* gene interact with the that of compatible *HD2* gene from an allelic pair of genes to generate active heterodimeric *HD1*-*HD2* transcription factor for achieving mating [41–42]. *A* MAT gene locus regulates the formation of clamp cell, dikaryon karyogamy, the synchronized division of dikaryon, the formation of the membrane between clamp cell and apical cell, and *B* MAT gene locus dominates the migration of nuclei, degradation of cell membranes and the cytomixis between clamp cell and secondary apical cell [32, 40, 43–44].

However, little is known about mating system and its association with virulence variation and high level of genetic diversity of *Pst* population since the discovery of its sexual cycle in 2010. Therefore, the objectives of the present study were to develop MAT genes of *Pst* and *Psh*, and validate the function of MAT genes to elucidate the mating system of *Pst* through selfing and hybridization of sexual reproduction. Furthermore, the study sought to investigate virulence variation and detecting MAT genes. It also aimed to assess the exchange and diversity of MAT genes to account for high genetic diversity in natural *Pst* populations in China.

## Results

### *HD* genes of *Pst* and *Psh*

A total of 13 *HD1* and *HD2* genes, comprising 11 from *Pst* and 2 from *Psh*, were cloned from the genomes of Chinese *Pst* and *Psh* isolates and subsequently sequenced. (S1 Fig; S1 Data.). Eleven *Pst HD1* genes (e series) and *HD2* genes (w series) included *PstHD1-e1/PstHD2-w1*, *PstHD1-e2/PstHD2-w2*, *PstHD1-e3/PstHD2-w3*, *PstHD1-e4/PstHD2-w4*, *PstHD1-e5/PstHD2-w5*, *PstHD1-e6/PstHD2-w6*, *PstHD1*-*e7/PstHD2-w7*, *PstHD1*-*e8/PstHD2*-*w8*, *PstHD1*-*e9/PstHD2*-*w9, PstHD1*-*e10/PstHD2*-*w10*, and *PstHD1*-*e11/PstHD2*-*w11* (S1A Fig). The two *Psh HD1* genes (e series) and *HD2* genes (w series) consisted of *PshHD1*-*e1/PshHD2*-*w1* and *PshHD1*-*e2/PshHD2*-*w2* (S1B Fig).

The analysis of physical arrangement *HD1-HD2* gene pair of the *Pst* and *Psh* showed that the sequences of PCR products amplificated by the primer 1 and 4 pair were same as those corresponding to *HD1* genes (S2 Fig). Likewise, the sequences of PCR amplicons produced with the primer 2 and 3 pair were identical to those corresponding to *HD2* genes (S2 Fig). These results indicated that the *HD1* and the *HD2* genes in *Pst* and *Psh* were linked (S1 and S2 Figs).

To further dissect the number of alleles of genes *HD1* and *HD2* in a single *P. striiformis* isolate, we performed statistics based on the NCBI Database (https://www.ncbi.nlm.nih.gov/). The number of *HD1* and *HD2* genes was calculated in *P. striiformis* genome on different hosts including wheat, barley, grass and triticale isolates (Table 1). Except for some *Pst* isolates due to low quality of genomic sequence, the number of *HD1* alleles or *HD2* alleles in the genomes of the dikaryotic *Pst* isolates was two, and those of the haploid *Pst* isolates was one. However, the number of *HD1* or *HD2* alleles in the genomes of *P*. *striiformis* isolates on barley and grass hosts were 0 and 1, respectively (Table 1).

**Table 1.**
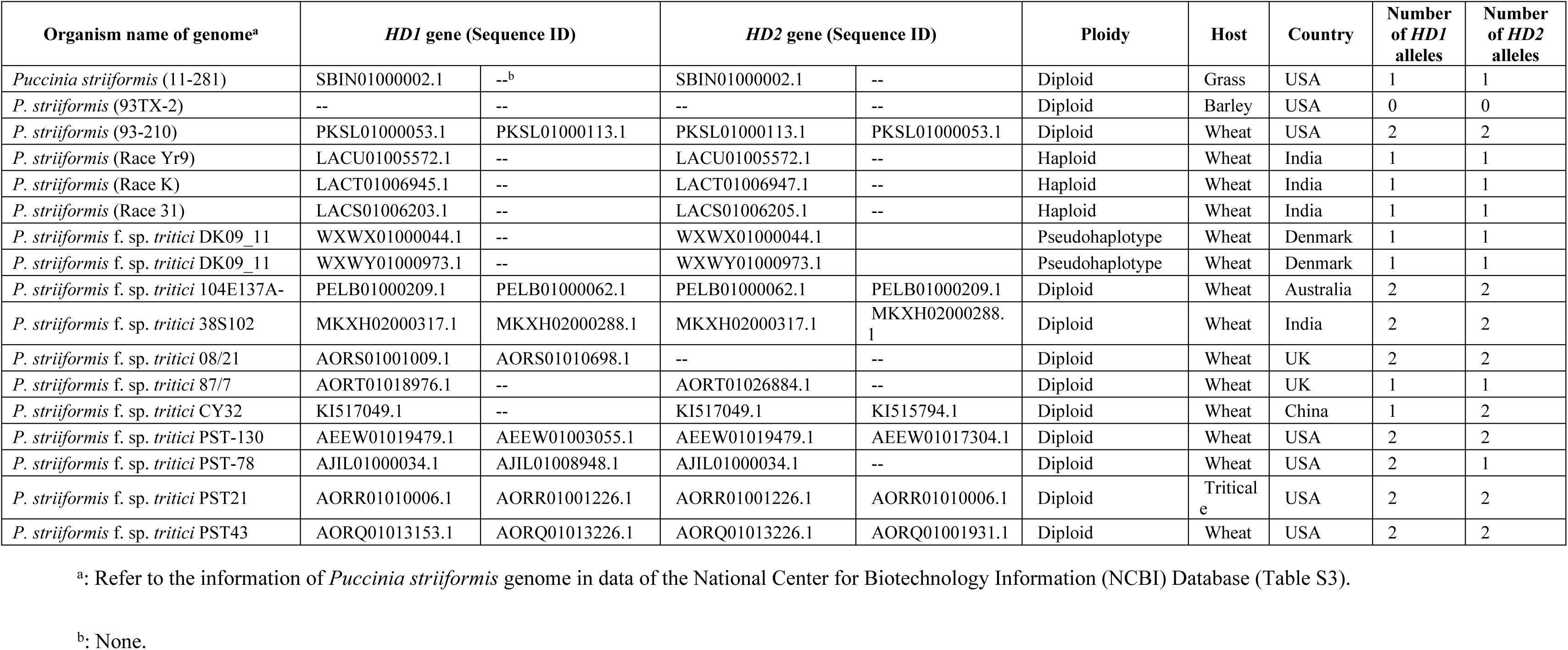
The number of *HD1* and *HD2* alleles in the genomes of *Puccinia striiformis* isolates released.

| Organism name of genome<sup>a</sup> | *HD1* gene (Sequence ID) |  | *HD2* gene (Sequence ID) |  | Ploidy | Host | Country | Number of *HD1* alleles | Number of *HD2* alleles |
| --- | --- | --- | --- | --- | --- | --- | --- | --- | --- |
| *Puccinia striiformis* (11-281) | SBIN01000002.1 | --<sup>b</sup> | SBIN01000002.1 | -- | Diploid | Grass | USA | 1 | 1 |
| *P. striiformis* (93TX-2) | -- | -- | -- | -- | Diploid | Barley | USA | 0 | 0 |
| *P. striiformis* (93-210) | PKSL01000053.1 | PKSL01000113.1 | PKSL01000113.1 | PKSL01000053.1 | Diploid | Wheat | USA | 2 | 2 |
| *P. striiformis* (Race Yr9) | LACU01005572.1 | -- | LACU01005572.1 | -- | Haploid | Wheat | India | 1 | 1 |
| *P. striiformis* (Race K) | LACT01006945.1 | -- | LACT01006947.1 | -- | Haploid | Wheat | India | 1 | 1 |
| *P. striiformis* (Race 31) | LACS01006203.1 | -- | LACS01006205.1 | -- | Haploid | Wheat | India | 1 | 1 |
| *P. striiformis* f. sp. *tritici* DK09_11 | WXWX01000044.1 | -- | WXWX01000044.1 |  | Pseudohaplotype | Wheat | Denmark | 1 | 1 |
| *P. striiformis* f. sp. *tritici* DK09_11 | WXWY01000973.1 | -- | WXWY01000973.1 |  | Pseudohaplotype | Wheat | Denmark | 1 | 1 |
| *P. striiformis* f. sp. *tritici* 104E137A- | PELB01000209.1 | PELB01000062.1 | PELB01000062.1 | PELB01000209.1 | Diploid | Wheat | Australia | 2 | 2 |
| *P. striiformis* f. sp. *tritici* 38S102 | MKXH02000317.1 | MKXH02000288.1 | MKXH02000317.1 | MKXH02000288.1 | Diploid | Wheat | India | 2 | 2 |
| *P. striiformis* f. sp. *tritici* 08/21 | AORS01001009.1 | AORS01010698.1 | -- | -- | Diploid | Wheat | UK | 2 | 2 |
| *P. striiformis* f. sp. *tritici* 87/7 | AORT01018976.1 | -- | AORT01026884.1 | -- | Diploid | Wheat | UK | 1 | 1 |
| *P. striiformis* f. sp. *tritici* CY32 | KI517049.1 | -- | KI517049.1 | KI515794.1 | Diploid | Wheat | China | 1 | 2 |
| *P. striiformis* f. sp. *tritici* PST-130 | AEEW01019479.1 | AEEW01003055.1 | AEEW01019479.1 | AEEW01017304.1 | Diploid | Wheat | USA | 2 | 2 |
| *P. striiformis* f. sp. *tritici* PST-78 | AJIL01000034.1 | AJIL01008948.1 | AJIL01000034.1 | -- | Diploid | Wheat | USA | 2 | 1 |
| *P. striiformis* f. sp. *tritici* PST21 | AORR01010006.1 | AORR01001226.1 | AORR01001226.1 | AORR01010006.1 | Diploid | Triticale | USA | 2 | 2 |
| *P. striiformis* f. sp. *tritici* PST43 | AORQ01013153.1 | AORQ01013226.1 | AORQ01013226.1 | AORQ01001931.1 | Diploid | Wheat | USA | 2 | 2 |

<sup>a</sup>: Refer to the information of *Puccinia striiformis* genome in data of the National Center for Biotechnology Information (NCBI) Database (Table S3).
<sup>b</sup>: None.

### Conserved HD homeodomain protein sequences

The HD1 and HD2 homeodomain protein sequences were derived from the translation of the cDNA sequences of the eleven *Pst HD1* and HD2 genes, as well as two *Psh* genes. The analysis showed that there were three helical regions (black block I, II, and III in red-dotted frames) in both the HD1 and HD2 homeodomains of *Pst* and *Psh* (S3A and 3B Figs). In the helix III block, the conserved amino acids of the DNA binding motif were WFXNXR, as reported in Basidiomycetes by Duboule [45]. Sequences alignment of HD1 and HD2 homeodomain protein of *Pst* and *Psh* displayed that sequence difference were mainly presented in the N-terminus rather than the C-terminus. Sequences of the HD2 homeodomain were more conserved than those of the HD1 (S3A and 3B Figs). The conserved DNA-binding domain sequences of the HD1 homology domains of 11 *Pst HD1* genes (*PstHD1-e1* to *PstHD1*-*e11*) and 2 *Psh HD1* genes (*PshHD1-e1* to *PshHD1*-*e2*) were WFRNAR, which was accordance with those in some species in Pucciniomycetes that were selected to use this study, but different from those of other ones (S1 Table and Fig 3C). Likewise, sequences of the conserved DNA-binding domain of the HD2 homology domains of 11 *Pst HD2* genes (*PstHD2-w1* to *PstHD2*-*w11*) and 2 *Psh HD2* genes (*PshHD2-w1* to *PshHD2*-*w2*) were WFCNAR, different from those of other species in Basidiomycetes that were selected to use in this study (S3D Fig).

Two phylogenetic trees were constructed based on the homeodomain protein sequences of *HD1* and *HD2* in *Pst* and *Psh*, as well as the selected species in Basidiomycetes. (Figs 1A and 1B). Each clustered was divided into five clades, including selected species of Microbotryomycetes (yellow shadow), Pucciniomycetes (orange shadow), Ustilaginomycetes (gray shadow), Tremellomycetes (green shadow), and Agaricomycetes (blue shadow). Homeodomain sequences of HD1 and HD2 of *Pst* and *Psh* isolates used in this study were involved in an individual subclade of Pucciniomycetes (Figs 1A and 1B). HD1 homeodomain sequences of *Pst* and *Psh* were closer with those of *P. graminis* f. sp. *tritici* (the causal pathogen of wheat stem rust, PgbE1, PgbE2) than *P. triticina* (the causal pathogen of wheat leaf rust, PtbE1) (Fig 1A). Whereas HD2 homeodomain sequences of *Pst* and *Psh* were more highly related to *P. triticina* (PtbEW1) than to *P. graminis* f. sp. *tritici* (PgbW1, PgbW2) (Fig 1B).

**Figure 1.**
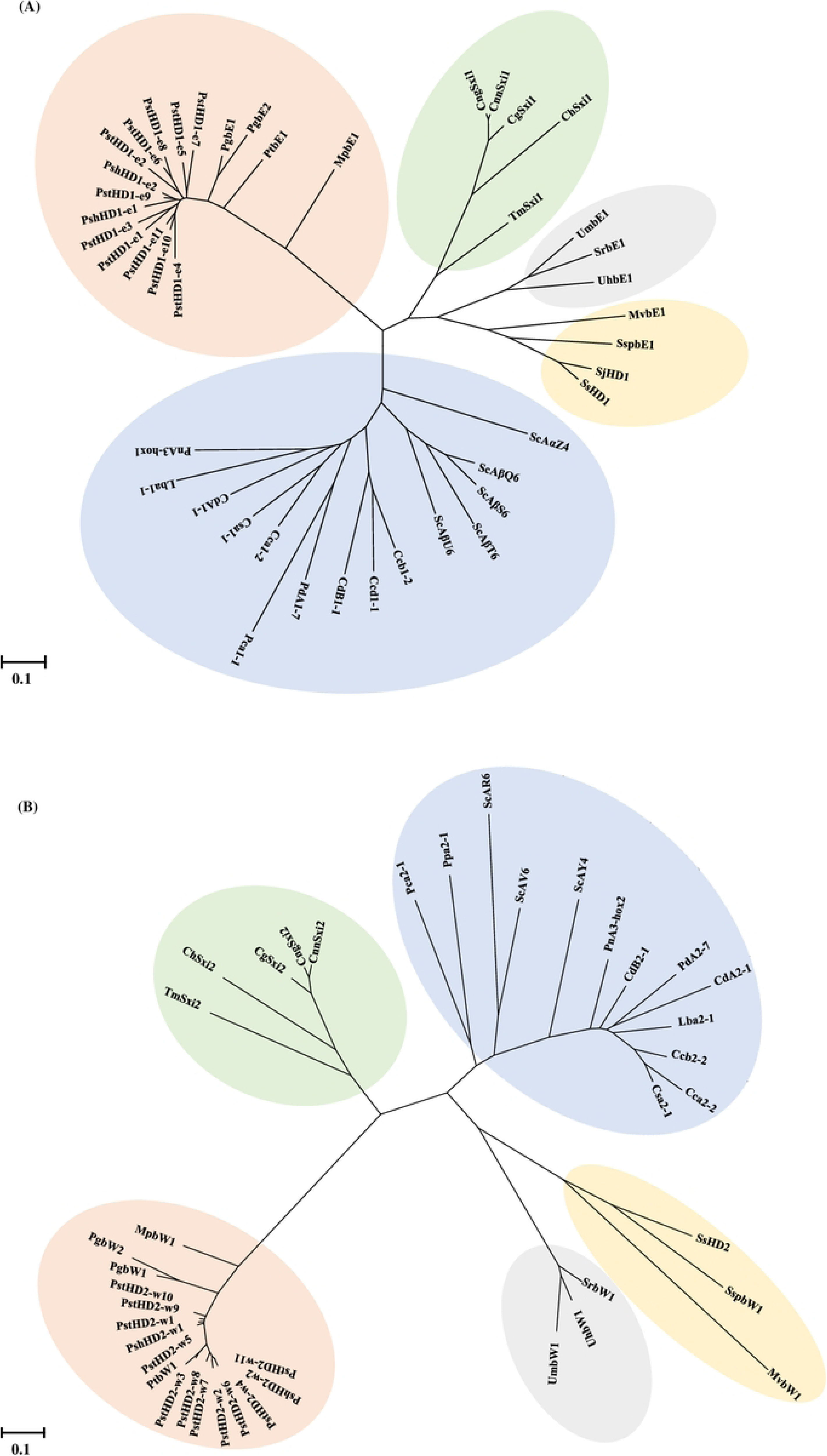
Phylogenetic trees showing different clusters of HD1 (a) and HD2 (b) genes of Puccinia striiformis f. sp. tritici, (PstHD1-e1 to PstHD1-e11, PstHD2-w1 to PstHD2-w11) and P. striiformis f. sp. hordei (PshHD1-e1 and PshHD1-e2, PshHD2-w1 and PshHD2-w2), and some fungi in Basidiomycetes listed in S1 Table.

### Mating system of *Pst*

During the pycnial stage of *Pst* race CYR32 on detached barberry leaves, 30 single-nectar (haploid pycniospore, n) were collected from pycnial lesions. It was utilized further to identify the types of *HD1* genes. Our result showed that 13 individual pycnium carried only *PstHD1-e1*, while 17 individual pycnium contained only *PstHD1-e10* (Fig 2A). The results indicating that *PstHD1-e1*of the *HD1* alleles located in a haploid pycniospore (n) with a mating type and *PstHD1-e10* of the *HD1* alleles in another one with opposite mating type separately. However, both *PstHD1-e1* and *PstHD1-e10* were not in an individual pycniospore simultaneously (Fig 2A).

**Figure 2.**
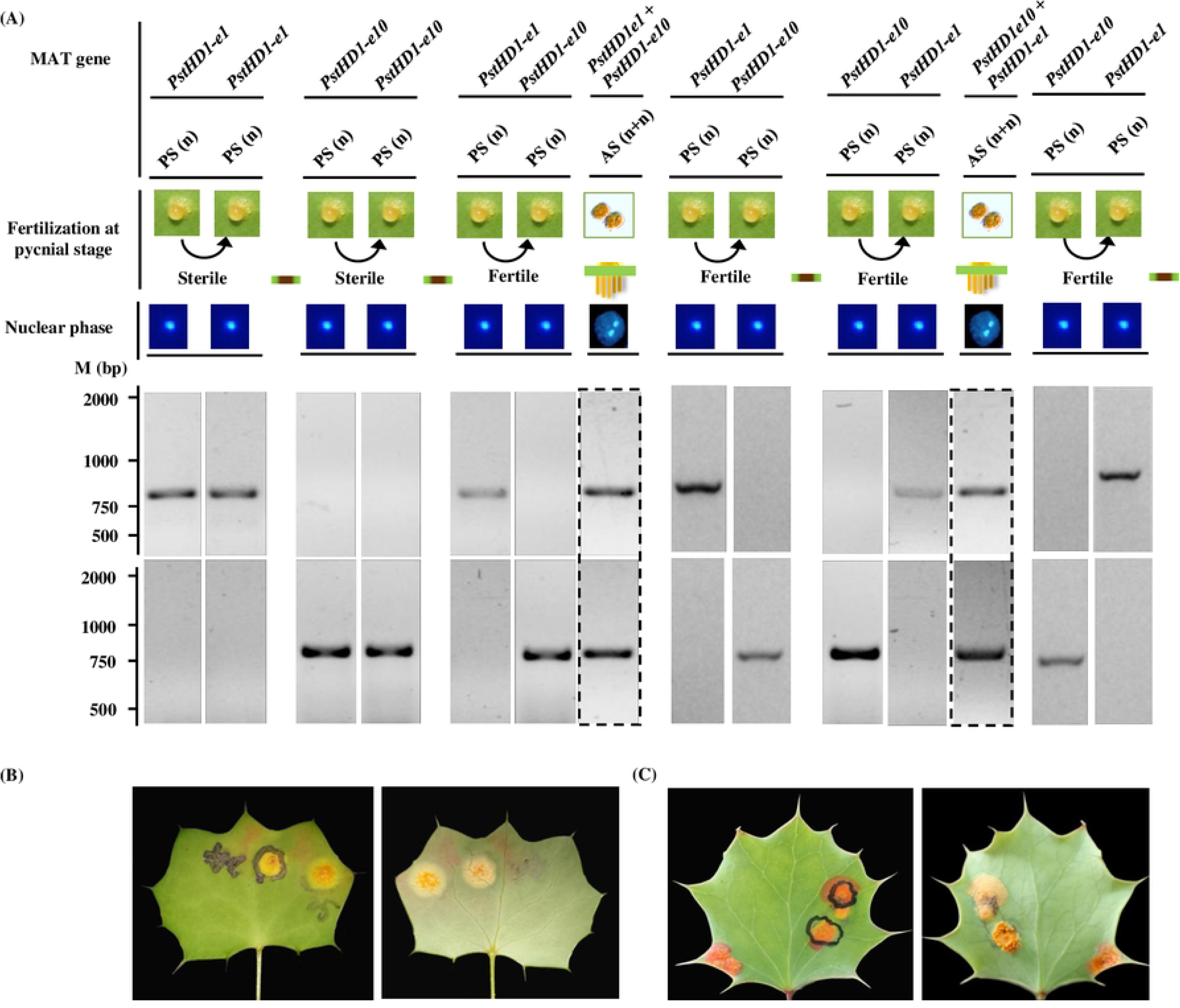
Aecia generation on barberry after mating between single-pycnia of *Puccinia striiformis* f. sp. *tritici* (*Pst*) race CYR32 by transferring one nectar of a pycnium to another. (a)Stained-agarose gels showing amplicons of pycniospores and selfed progeny’s urediniospores of *Pst* race CYR32 amplified with *HD1* gene-specific primers based on PCR amplification with *PstHD1-e1* and *PstHD1-e10* gene-specific primers, respectively. M, DL 2000 DNA Marker (Shanghai Yuanye Bio-Technology Co., Ltd., China). PS = Pycniospores. AS = Aeciospores. (b)A experimental example showing no aecia generation on detached-leaves of barberry (*Berberis shensiana*) after mating between two single pycnium with same *PstHD1* genes (*PstHD1-e1* or *PstHD1-e10*). (c) Aecia produced on an individual pycnium lesion after fusion between one pycnium with *PstHD1*-*e1* or *PstHD1*-*e10* gene and the other with *PstHD1-e10* or *PstHD1-e1* gene.

Regardless of fusions between two nectars carrying *PstHD1-e1* or between two ones carrying *PstHD1-e10* on detached barberry leaves at the pycnial stage, no aecia were observed to develop on pycnia lesions one week or longer after fusion (Figs 2A and 2B). In contrast, eleven of the fusions between single nectar containing *PstHD1*-*e1* and that carrying *PstHD1*-*e10* by orthogonal and reverse ways resulted in the production of five aecia. The mating rate ranged from 33.3% to 60.0%, with an average of 45.5% (Fig 2C and Table 2). Molecular detection of *HD1* genes with primers specific *PstHD1*-*e1* and *PstHD1*-*e10* among selfed progenies showed that both *PstHD1*-*e1* and *PstHD1*-*e10* were detected in each of progenies, indicating that the two *HD1* genes from two different pycniospores recombined into an individual progeny through fusions (Fig 2A).

**Table 2.** Production of aecia by mating between pycnia of *Puccinia striiformis* f. sp. *tritici* race CYR32.

| Type | Way of mating<sup>a</sup> | No. of mating | No. of pycnia generating aecia | No. of pycnia without generating aecia | Percentage of producing aecia (%) | | <sup>b</sup>Confidence level ($P < 100\%$) |
| --- | --- | --- | --- | --- | --- | --- | --- |
| I | e1→e1 | 9 | 0 | 3 | 0.0 | 0.0 | -- |
|  | e10→e10 |  | 0 | 6 | 0.0 |  |  |
| II | e1→e10 | 11 | 3 | 2 | 60.0 | 45.5 | 99.5 |
|  | e10→e1 |  | 2 | 4 | 33.3 |  |  |

<sup>a</sup> E1→e1, the mating between pycnia containing *PstHDI-e1*; e10→e10, mating between pycnia containing *PstHDI-e10*; e1→e10, the nectar from one pycnium containing *PstHDI-e1* was smeared to another pycnium containing *PstHDI-e10* for mating; e10→e1, the nectar from one pycnium containing *PstHDI-e10* was smeared to another pycnium containing *PstHDI-e1* for mating.
<sup>b</sup> the confidence level with the probability of producing aecia of less than 100% based on interval estimation of 0-1 distribution parameter.

Based on the 0-1 distribution parameter, after mating between pycnia with different *HD1* gene types, the confidence level with the probability (*P*) of less than 100% of producing aecia was tested. The results in this study indicated that after mating, confidence interval was 99.5%, namely the probability of producing aecia is less than 100%. Thus, other loci unlinked with *HD* locus in *Pst*, hinting to play biological functions in sexual recombination. Accordingly, the mating system of *Pst* is tetrapolar.

### *HD* gene composition and virulence phenotype of *Pst* progenies via selfing and crossing

Ten selfed progenies were obtained from the *Pst* race CYR32 (S_1_-1 to S_1_-10), and five were derived through crossing between CYR32 and CYR23 (F_1_-1 to F_1_-5; Table 3). The *HD* gene composition of the parental races (CYR23 and CYR32), selfed progenies of CYR32, and crossed progenies of CYR23 and CYR32 were subsequently detected (Table 3 and S4 Fig). Detections of the *HD1* and *HD2* genes revealed that race CYR32 harbored combinations of *PstHD1*-*e1* and *PstHD1*-*e10*, as well as *PstHD2*-*w1* and *PstHD2*-*w10*. Similarly, race CYR23 exhibited combinations of *PstHD1*-*e3* and *PstHD1*-*e8*, and *PstHD2*-*w3* and *PstHD2*-*w8* (Table 3). Selfed progenies exhibited the same combination types of the *HD1* and of *HD2* genes as the parental races (Table 3). While the crossed progenies between CYR23 and CYR32 races showed gene exchange and recombination, and exhibited two new combination types of *HD1* and *HD2* genes. One was *PstHD1*-*e1* and *PstHD1*-*e8* (*PstHD2*-*w1* and *PstHD2*-*w8*), and the other was *PstHD1*-*e3* and *PstHD1*-*e10* (*PstHD2*-*w3* and *PstHD2*-*w10*) (Table 3). It reflected interspecific sexual hybridization is an important way to form new recombination types.

**Table 3.**
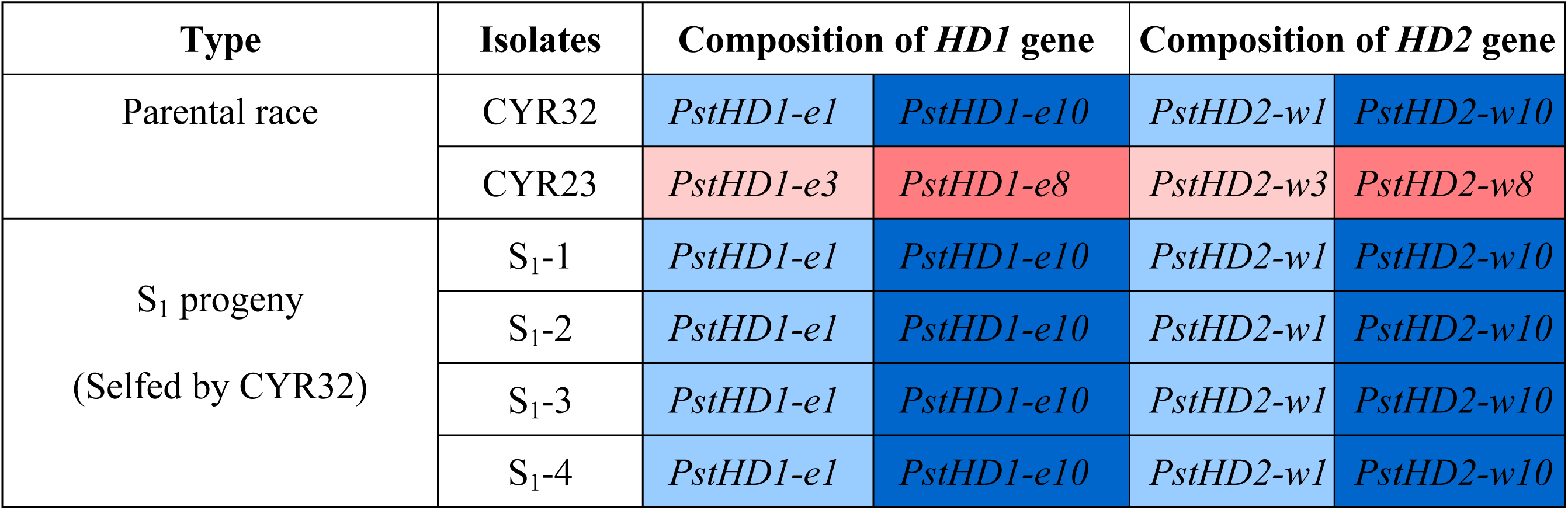
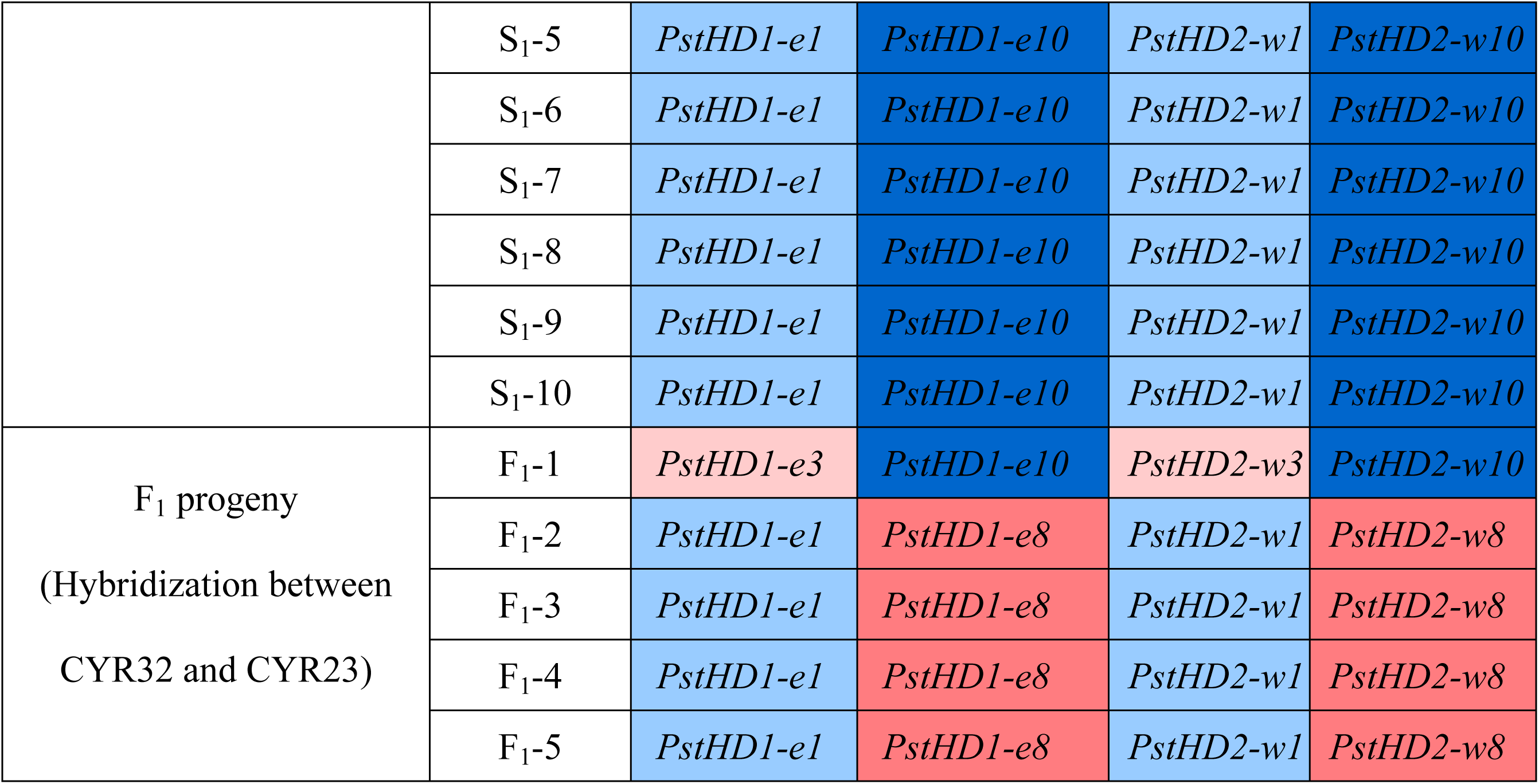
Composition of mating type genes *HD1* and *HD2* of parent and progeny isolates of *Puccinia striiformis* f. sp. *tritici*.

| Type | Isolates | Composition of *HD1* gene |  | Composition of *HD2* gene |  |
| --- | --- | --- | --- | --- | --- |
| Parental race | CYR32 | *PstHD1-e1* | *PstHD1-e10* | *PstHD2-w1* | *PstHD2-w10* |
|  | CYR23 | *PstHD1-e3* | *PstHD1-e8* | *PstHD2-w3* | *PstHD2-w8* |
| S<sub>1</sub> progeny<br>(Selfed by CYR32) | S<sub>1</sub>-1 | *PstHD1-e1* | *PstHD1-e10* | *PstHD2-w1* | *PstHD2-w10* |
|  | S<sub>1</sub>-2 | *PstHD1-e1* | *PstHD1-e10* | *PstHD2-w1* | *PstHD2-w10* |
|  | S<sub>1</sub>-3 | *PstHD1-e1* | *PstHD1-e10* | *PstHD2-w1* | *PstHD2-w10* |
|  | S<sub>1</sub>-4 | *PstHD1-e1* | *PstHD1-e10* | *PstHD2-w1* | *PstHD2-w10* |
|  | S<sub>1</sub>-5 | *PstHD1-e1* | *PstHD1-e10* | *PstHD2-w1* | *PstHD2-w10* |
|  | S<sub>1</sub>-6 | *PstHD1-e1* | *PstHD1-e10* | *PstHD2-w1* | *PstHD2-w10* |
|  | S<sub>1</sub>-7 | *PstHD1-e1* | *PstHD1-e10* | *PstHD2-w1* | *PstHD2-w10* |
|  | S<sub>1</sub>-8 | *PstHD1-e1* | *PstHD1-e10* | *PstHD2-w1* | *PstHD2-w10* |
|  | S<sub>1</sub>-9 | *PstHD1-e1* | *PstHD1-e10* | *PstHD2-w1* | *PstHD2-w10* |
|  | S<sub>1</sub>-10 | *PstHD1-e1* | *PstHD1-e10* | *PstHD2-w1* | *PstHD2-w10* |
| F<sub>1</sub> progeny<br>(Hybridization between<br>CYR32 and CYR23) | F<sub>1</sub>-1 | *PstHD1-e3* | *PstHD1-e10* | *PstHD2-w3* | *PstHD2-w10* |
|  | F<sub>1</sub>-2 | *PstHD1-e1* | *PstHD1-e8* | *PstHD2-w1* | *PstHD2-w8* |
|  | F<sub>1</sub>-3 | *PstHD1-e1* | *PstHD1-e8* | *PstHD2-w1* | *PstHD2-w8* |
|  | F<sub>1</sub>-4 | *PstHD1-e1* | *PstHD1-e8* | *PstHD2-w1* | *PstHD2-w8* |
|  | F<sub>1</sub>-5 | *PstHD1-e1* | *PstHD1-e8* | *PstHD2-w1* | *PstHD2-w8* |

A total of 15 *Pst* isolates, including 2 parental races, 10 selfed progenies and 3 crossed progenies (hybrids), were tested for virulence phenotypes (VP). The results showed that virulence formula of 13 selfed and crossed progenies was obviously distinguished from that of the two parental races (VP4 and VP15). Within crossed progenies, those of which, with the same combination type of *HD1* or *HD2* gene, exhibited different virulence phenotypes, such as F_1_-2 and F_1_-3 (Table 4).

**Table 4.**
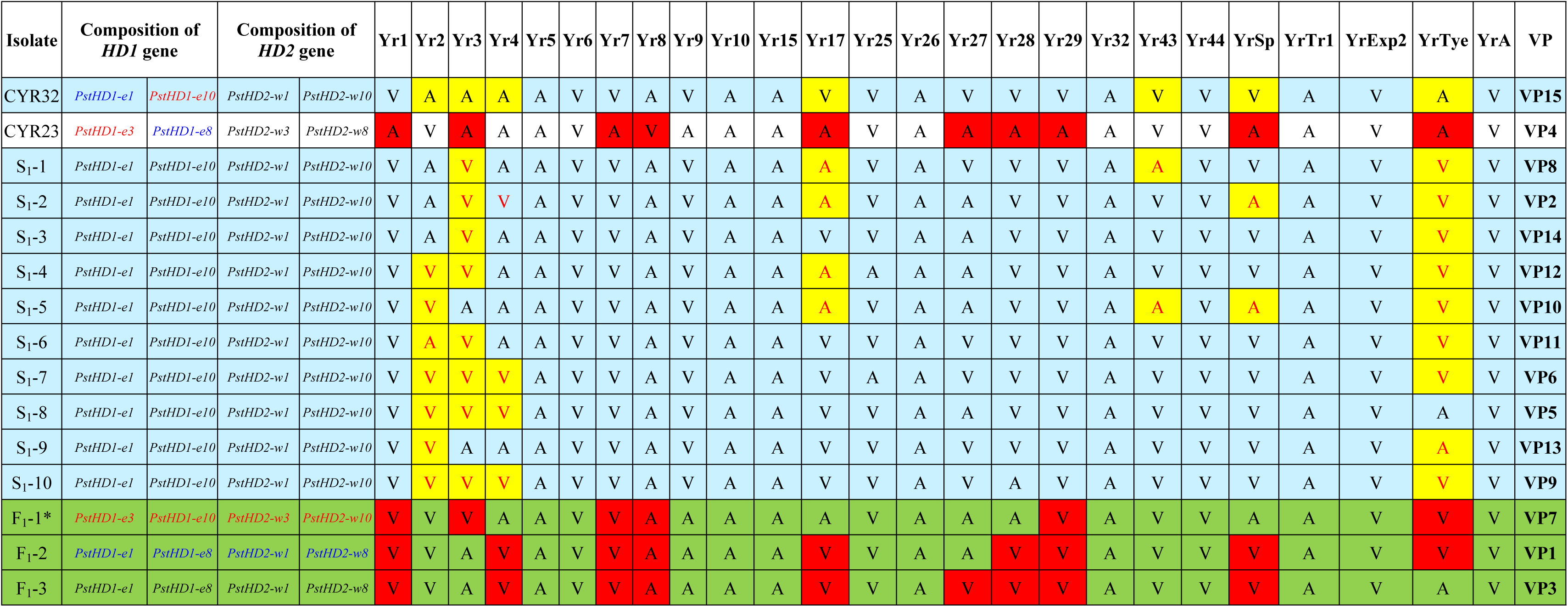
Virulence phenotypes (VPs) of the constructed sexual population of *Puccinia striiformis* f. sp. *tritici* on 25 wheat *Yr* gene lines.

### Detection, geographic distribution and proportion of *HD* genes in Chinese natural *Pst* population

Totally, 213 stripe rust (*Puccinia striiformis*, *Ps*) isolates, collected from 92 sampling sites in 11 provinces of China, including 200 *Pst* isolates from Tibet (19), Xinjiang (16), Qinghai (16), Gansu (31), Shaanxi (18), Sichuan (16), Yunnan (15), Guizhou (17), Henan (23), Hubei (14), and Jiangsu (15), and 13 *Psh* isolates from Tibet, were used in this study (S2 Table). Overall 213 *Ps* isolates were tested by using 13 pairs of *HD1* specific-primers and 13 pairs of *HD2* specific-primers (S3 and S4 Tables). The testing results of *HD* gene of 213 *Ps* isolates samples are shown in the Fig 3, in which the testing results of *HD1* gene and *HD2* gene of each sample correspond to Figs 3A and 3B, respectively. We can find that the linked *HD1* gene and *HD2* gene type both appear in the same sample of natural *Ps* population at the same time (Fig 3C). Therefore, the following analysis, such as *HD* gene distribution and proportion and *HD* gene-based population structure, only needs to be based on either the *HD1* gene or the *HD2* gene.

**Figure 3.**
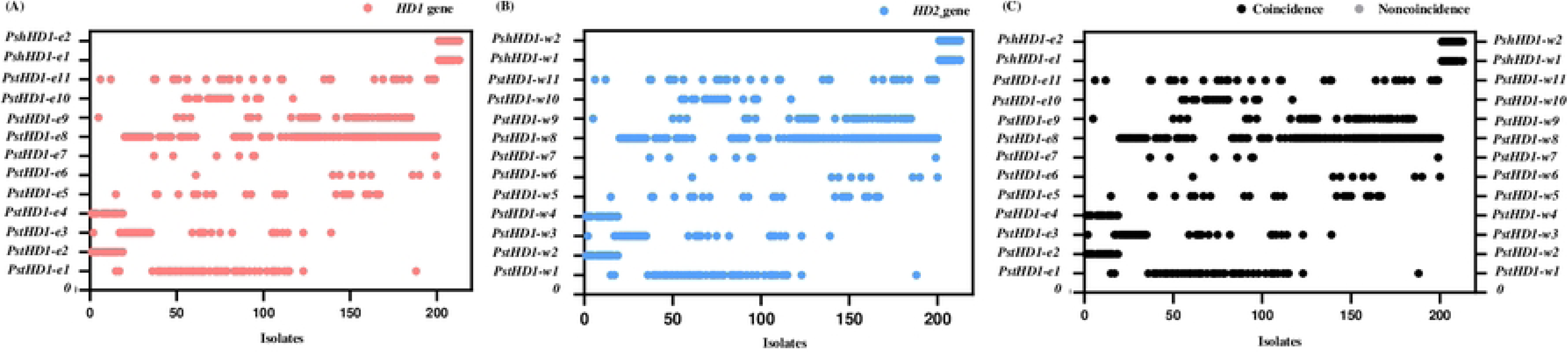
Distribution and coincidence analysis of *HD1* and *HD2* genes of 213 isolates of a natural *Puccinia striiformis* population from 11 provinces in China. Numbers from 1 to 200 indicated *Puccinia striiformis* f. sp. *tritici* (*Pst*) isolates collected from wheat, and those from 201 to 213 indicated *Puccinia striiformis* f. sp. *hordei* (*Psh*) isolates collected from barley. (a) *HD1* gene. (b) *HD2* gene. (c) Coincidence of *HD1* and *HD2* genes.

Among the tested sequences, 16 exhibited mutations. Of these, 14 had a mutation rate less than 1%, while two had a rate exceeding 1%. The highest mutation rate was 2.78% (S5 Table). The *Pst* isolates harboring mutations in the *HD* gene were mainly distributed in Sichuan province. Notably, the *Pst* isolate with the highest mutation rate in the *HD1* gene was also sourced from Sichuan province. Based on the test results, the distribution of the *HD1* gene of *Ps* in 11 provinces of China was statistically analyzed (Fig 4A). The gene *PshHD1*-*e1* and *PshHD1*-*e2* (cloned from the *Psh* genome) was only detected in *Psh* isolates, and none of the *Pst* isolates collected in 11 provinces (including Tibet) were detected this gene, which indicated the gene is unique to *Psh*. The genes *PstHD1*-*e2* and *PstHD1*-*e4* were only distributed in the *Pst* population of Tibet. In contrast, the *PstHD1-e8* gene exhibited the most extensive distribution, being present in *Pst* populations across ten provinces except Tibet, followed by *PstHD1*-*e11* and *PstHD1*-*e9*, which present in *Pst* populations in 9 and 8 provinces, respectively.

**Figure 4.**
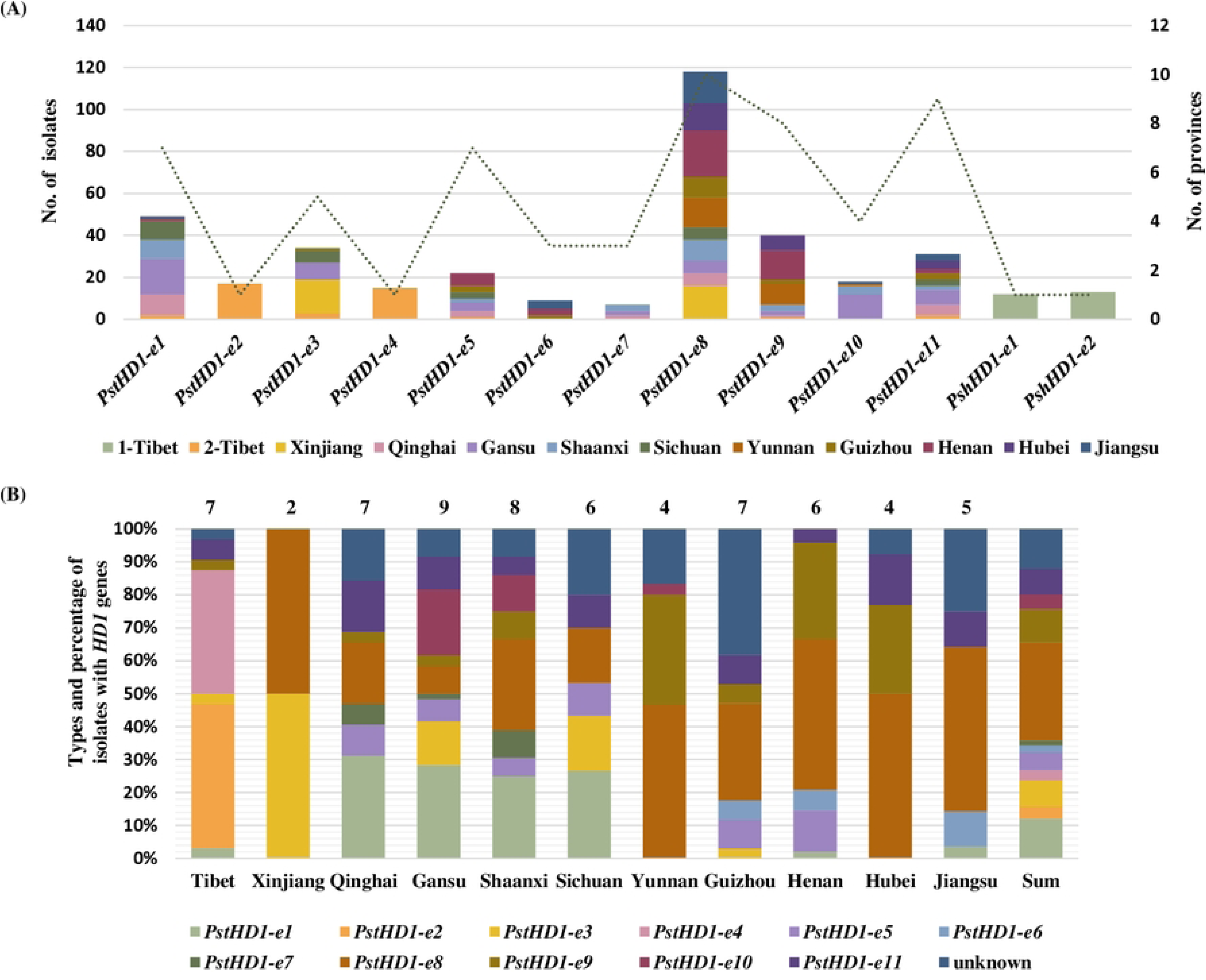
Distribution and percentage of *HD1* gene in 213 isolates of a natural *Puccinia striiformis* population from 11 provinces in China. (a) Distribution of *HD1* gene. 1-Tibet and 2-Tibet indicated that isolates of *Puccinia striiformis* f. sp. *hordei* (*Psh*) from barley, and that those of *Puccinia striiformis* f. sp. *tritici* (*Pst*) from wheat, respectively. (b) Percentage of *HD1* gene in each of 11 provinces. “Unknown” types indicated that the *HD1* gene types designed in this study was not detected among isolates in which new *HD1* gene type or the *HD1* allele deletion are possibly contained.

Based on the number of *HD1* alleles in the single-urediniospore *Pst* isolate genome is 2 (Table 1), the proportion of *HD1* gene contained in *Pst* isolates collected from 11 provinces in China was statistically analyzed (Fig 4B). The gene *PstHD1-e8* held the most significant proportion, comprising 30% and it occupied a dominant position in the *Pst* populations in nine provinces: Xinjiang, Qinghai, Sichuan, Yunnan, Shaanxi, Guizhou, Hubei, Henan and Jiangsu. The “unknown” genes (new *HD1* gene types or *HD1* gene deletions), accounted for 12%. *PstHD1*-*e1* exhibited dominance in Qinghai, Gansu, Shaanxi, and Sichuan provinces, while *PstHD1*-*e9* dominated in Yunnan, Henan, and Hubei provinces. The dominant *HD1* genes in Tibet were *PstHD1*-*e2* and *PstHD1*-*e4*. A comparative analysis of the *HD1* gene types in populations across eleven provinces revealed that Xinjiang had the least types, with only *PstHD1-e3* and *PstHD1-e8*. Gansu and Shaanxi provinces, on the other hand, had relatively more types, with at least eight and seven types, respectively. To further take an insight into the frequency of gene recombination in *Pst* natural populations across various provinces or regions of China, we conducted statistics and analysis of the *HD* gene combination types in Chinese natural populations (Fig 5 and S6 Table). There were at least 30 *HD* gene combination types in the *Pst* populations in 11 provinces of China, among which 6 combination types were composed of “unknown” genes. The *HD* gene combination types of *Pst* populations in Gansu and Shaanxi provinces were more abundant than those in other provinces. Gansu province contained at least 16 types, and Shaanxi province contained at least 10 types (Fig 5A). The *HD1* gene combination type of *Pst* population in Xinjiang province was relatively single, with only one combination type T4 (Fig 5A). Moreover, the *Pst* populations in other provinces except Xinjiang all had more than 2 types of *HD* gene combinations.

**Figure 5.**
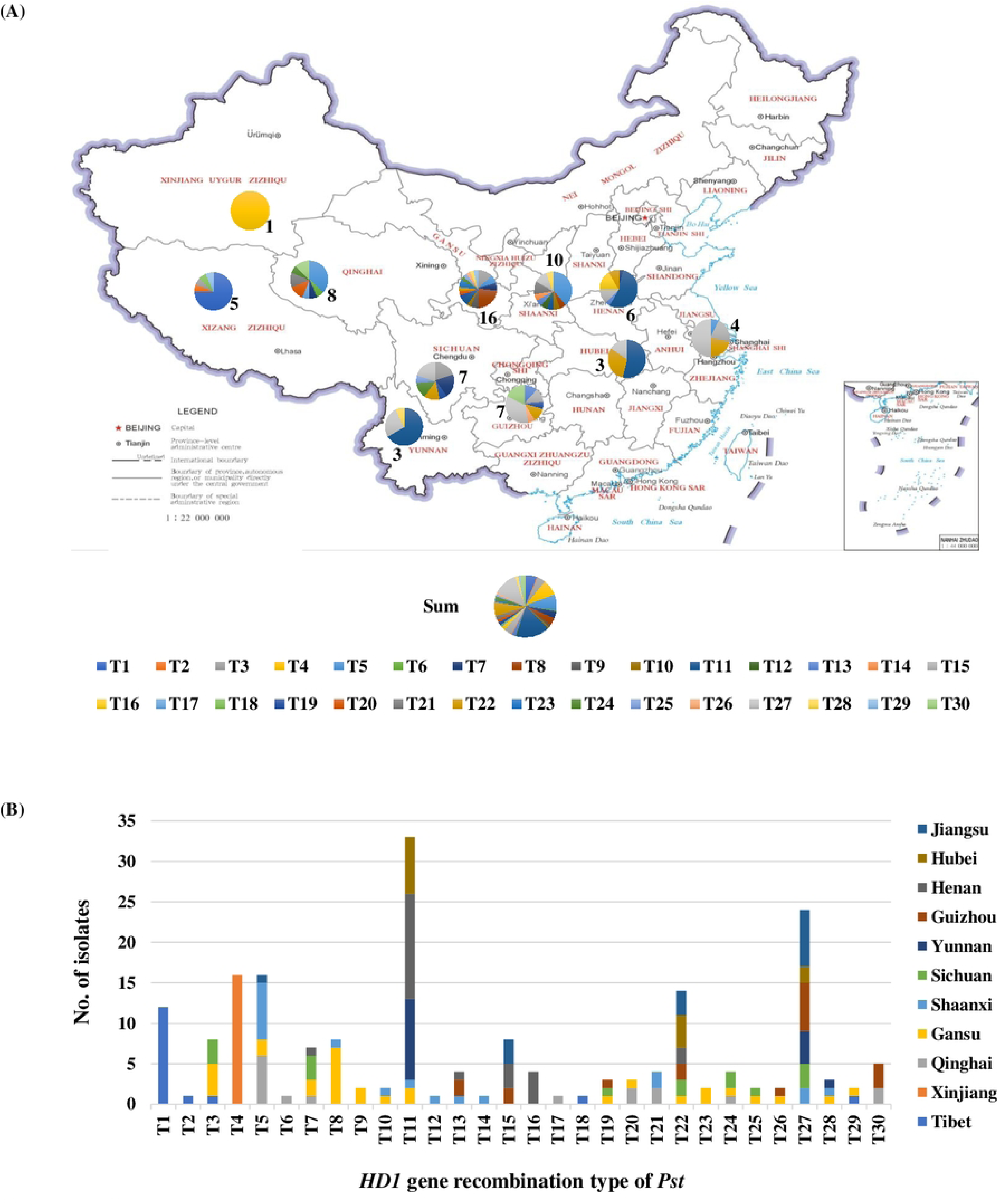
Distribution and percentage of *HD1* gene combination types of *Puccinia striiformis* f. sp. *tritici* isolates in each of 11 provinces in China. (a) The Distribution of *HD1* gene combination types in each of 11 provinces in China. (b) Number and percentage of *HD1* gene combination types in each of 11 provinces in China. The *HD1* gene combination types listed in S6 Table.

The *Pst* with *HD* gene combination type T1 (*PstHD1*-*e2* and *PstHD1*-*e4*, *PstHD2*-*w2* and *PstHD2*-*w4*) was only found in Tibet but not in other provincial populations, and this type of isolates were dominant in Tibet (Fig 5B). The *Pst* with type T4 (*PstHD1*-*e3* and *PstHD1*-*e8*, *PstHD2*-*w3* and *PstHD2*-*w8,*) was only found in Xinjiang, and this type of isolates were dominant in Xinjiang (Fig 5B).

### Population structure and cluster analysis of natural *Pst* population

Based on the test result of the *HD1* gene (under the background of genome in dikaryotic *Pst* containing 2 *HD1* alleles), 194 *Pst* isolates from 11 provinces were conducted for population structure and cluster analysis by using STRUCTURE software, Discriminant Analysis of Principal Components (DAPC) and software POPGENE.

Based on STRUCTURE analysis, we get the optimum cluster level at *K*=8, different population genetic cluster groups were showed with different color in bar chart (Fig 6A). The blue cluster was dominated in Tibet, the green cluster was dominated in Xinjiang province, and the other clusters in the two provinces had a lower proportion. Gansu and Sichuan provinces were dominated by the orange cluster. Qinghai province and Shaanxi province were dominated by red cluster. Yunnan, Henan and Hubei provinces were dominated by the gray cluster. The gray, and white clusters of Jiangsu province were dominant. Guizhou province was dominated by white, yellow, purple, and green clusters. The *Pst* population in Tibet and Xinjiang provinces both had unique cluster composition.

**Figure 6.**
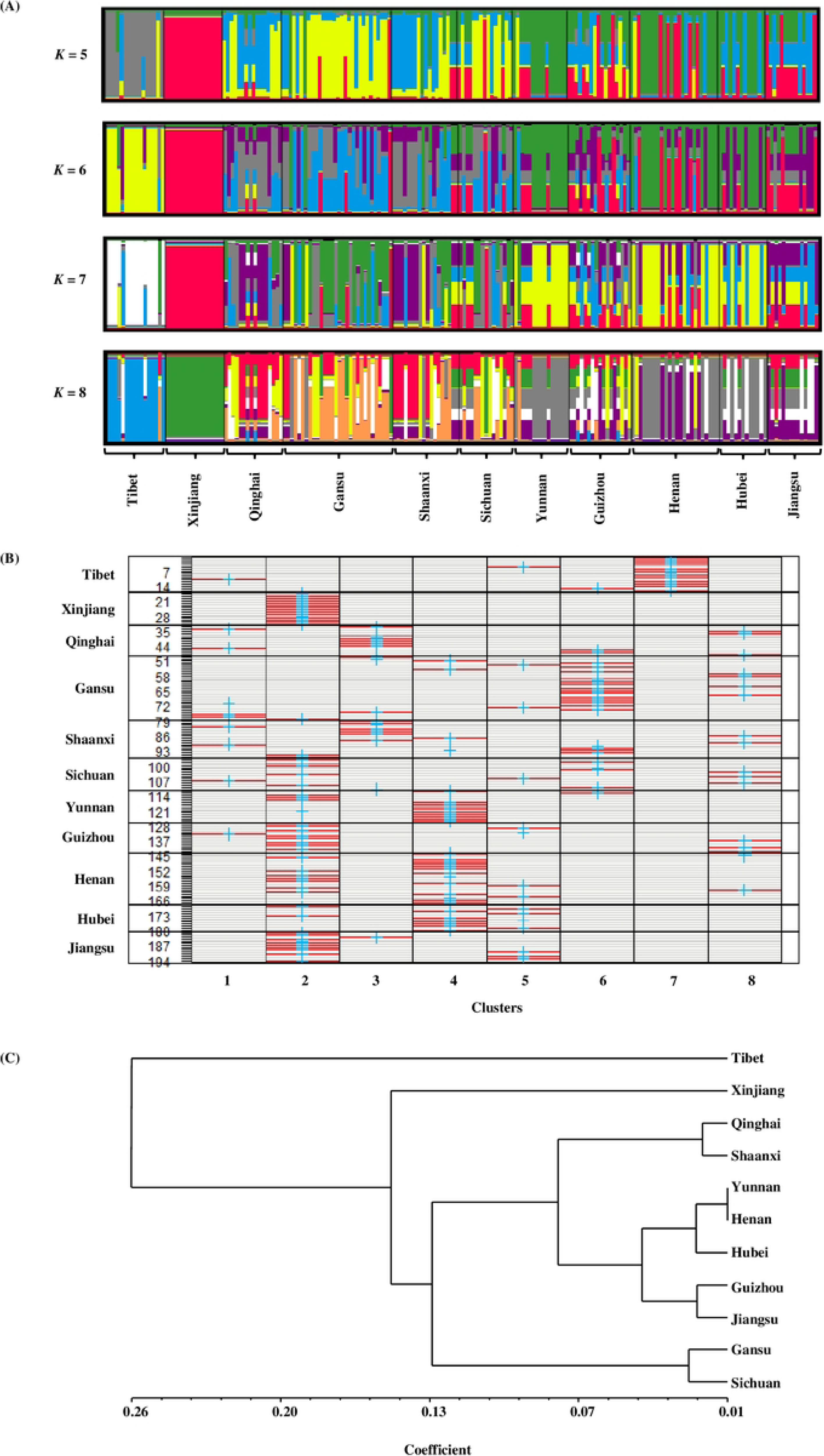
Population structure and cluster analysis of *Puccinia striiformis* f. sp. *tritici* (*Pst*) isolates from 11 provinces of China based on *HD1* gene. (a)Population structure of *Pst* isolates from 11 provinces of China based on *HD1* gene under the *K* vales ranging from 5 to 8 (optimal *K =* 8). (b) Assignation plot based on the discriminant analysis of principal components (DAPC) performed on *HD1* genes of *Pst* isolates from 11 provinces of China. Heat colors represented membership probabilities (red = 1, white = 0). Blue crosses represented the prior cluster provided to DAPC, and those of which on red clusters indicated DAPC classification of isolates was consistent with the original clusters, while the remaining of which were not on red clusters may be inferred admixed or misclassified. (c) Dendrogram based on genetic identity of *HD1* genes of the clusters of *Pst* isolates from 11 provinces of China.

Population genetic structure was further validated through DAPC analysis, which also divided the overall China population of 11 provinces into 8 cluster groups (S5 Fig and Fig 6B). This result showed that Cluster 7 was relatively independent from other Clusters, and Cluster 7 was only composed of *Pst* population from Tibet, indicating that the *Pst* population in Tibet is a relatively independent population (Fig 6B). It also found the trace of cluster 2 in Xinjiang region, which was composed of multiple local *Pst* population (Fig 6B). It indicated that the *Pst* population of Xinjiang may have some exchanges with those of neighbor provinces. Whereas the results were basically consistent with the results of STRUCTURE analysis.

The software POPGENE (Version 1.32) was used to calculate the genetic identity of *Pst* isolates from 11 provinces in China. Based on the genetic identity of the *Pst* populations in each province, a dendrogram was constructed by the UPGMA method (Fig 6C). The dendrogram indicated that overall isolates from 11 provinces were clustered into 5 clades. Each of the *Pst* populations from both Tibet and Xinjiang provinces corresponded to respective independent clade distinguished from other clades. The *Pst* populations from Gansu and Sichuan provinces clustered in a same cluster. The *Pst* populations of Qinghai and Shaanxi provinces were clustered into a subclade. Five *Pst* populations including Yunnan, Henan, Hubei, Guizhou and Jiangsu provinces were clustered into a subclade.

### Virulence phenotypes of natural *Pst* population

A total of 24 *Pst* isolates, selected from Gansu, Guizhou, Henan, Shaanxi, Tibet, Xinjiang and Yunnan provinces, tested for *HD1* gene and virulence phenotypes (VPs) on 25 wheat lines carrying different single *Yr* genes. The 24 isolates were identified as 21 various VPs, VP1-VP21, and included 9 different *HD1* gene combination types (Table 5).

**Table 5.**
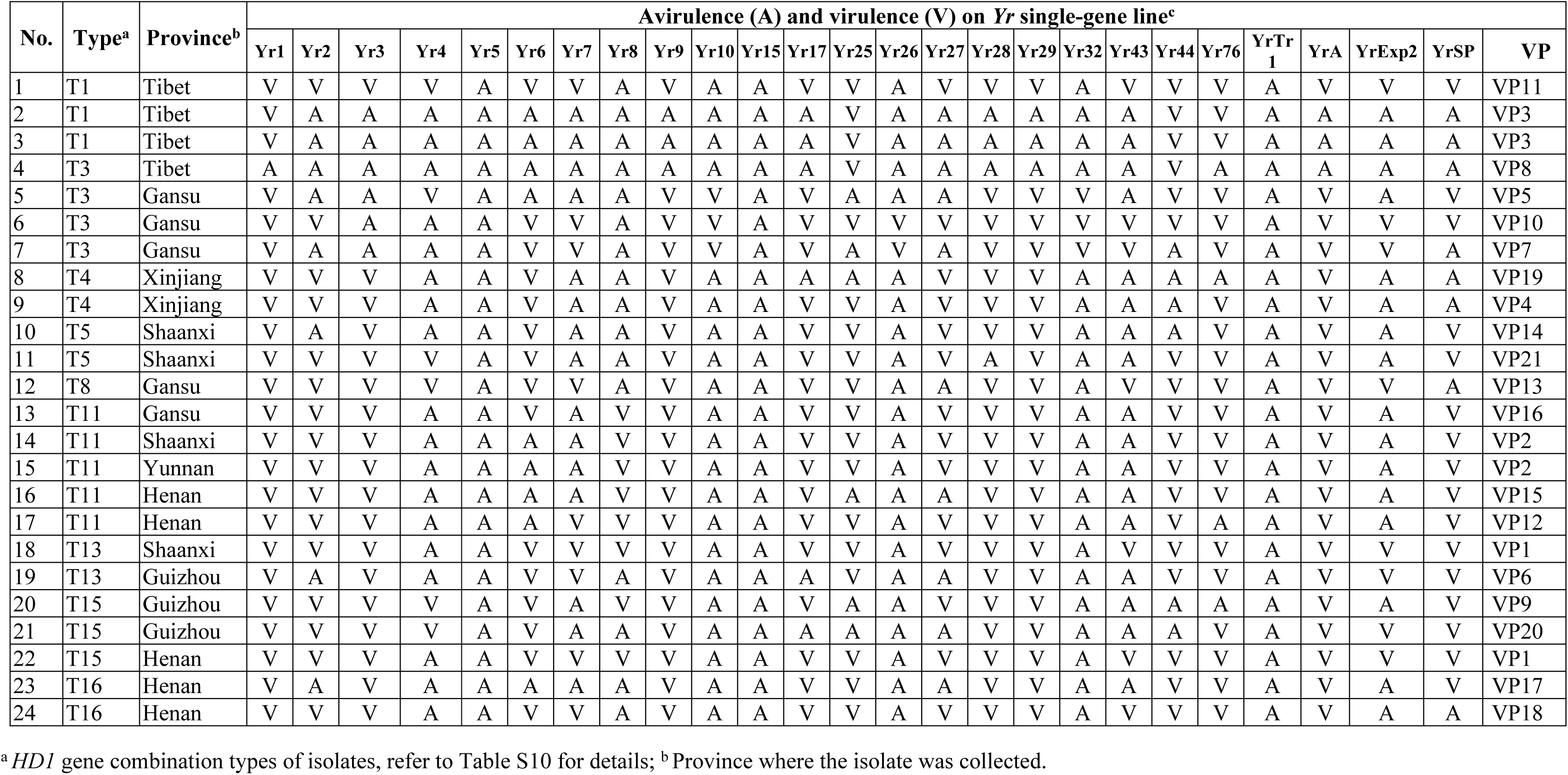
Virulence phenotypes (VPs) of the selected *Puccinia striiformis* f. sp. *tritici* isolates collected in China on 25 wheat *Yr* gene lines.

Among the 24 *Pst* isolates, most of which contained different *HD1* gene combination types exhibited various VPs (Table 5). Whereas, some isolates with different *HD1* gene combinations can be detected the same VPs. It was exemplified by the isolates (n = 18), belonging to type T13, had the same VP formula as the isolate (n = 22) that was categorized in type T15. However, most of these isolates contained same *HD1* gene combination types exhibited various VPs (Table 5). It was exemplified by the isolate (n= 1), shared the same *HD1* gene combination type T1 with the isolates (n = 2), but exhibited different VP patterns, respectively.

## Discussion

### Evolutionary considerations of mating type of *Pst*

The presence of all known types of genes (*HD* and *P/R*) with mating type functions is remarkably ancient, suggesting that they may have originated once in the most recent common ancestor of Basidiomycetes. However, it is possible that Basidiomycetes may have multiple independent origins [36, 46]. In this study, we constructed a phylogenetic tree based on the HD1 and HD2 homeodomain sequences of basidiomycetes (Fig 1). Result reflected species belonging to the same order in taxon were grouped in the same cluster. *Puccinia striiformis* was placed in the cluster of Pucciniomycetes and was closer to *P. graminis* and *P. triticina* than to *Melampsora laricis-populina* in terms of genetic relationship. This genetic association was in accordance with the biological evolution of Basidiomycetes. Those results suggest that the degree of evolution of the *HD* gene in Basidiomycota is generally consistent with its biological evolution, indicating that the HD homeodomain was certain conserved.

In this study, 13 *HD1* alleles were discovered from natural *Ps* populations with highly divergent sequences. Forming such differential sequences is a formidable process due to evolutionary limitations [36]. However, due to the compatibility and functionality of *HD* genes, ensuring that a pair of the *HD1* and *HD2* partner genes are compatible and functional, deletion or loss of function of the remaining *HD1* gene or *HD2* gene does not to be invalid of sexual fertility [39–40]. In the natural *Pst* population, there were combinations of one *HD1* gene (or *HD2* gene) with many different type *HD1* genes (or *HD2* genes), and most of the *HD1* genes (or *HD2* genes) had this phenomenon (S7 Table). This result confirmed that *HD1* genes (or *HD2* genes) had certain compatibility. In view of an evolutionary perspective, the reason for the presence of many *HD1* and *HD2* gene types with significant sequence differences in the natural *Ps* population could be explained that *HD* genes had compatibility. Whereas the remaining *HD1* or *HD2* genes, even if mutated, would have relatively little impact on the function of HD transcription factors, and the pressure on them by natural selection would be reduced. Thus, genetic variation could inherit and accumulate. According to the analysis of the constructed phylogenetic tree, we found that the degree of evolution between linked *HD1* and *HD2* gene was not consistent (Fig 1). This result consistent with the above speculation.

The role of recombination and mutations in generating new MAT alleles is not clear from current studies. In this study, mutations were detected in some types of *HD1* genes in natural *Pst* populations, with a mutation rate of up to 2.78%, and the mutation rate of *HD2* gene was relatively low (0.40%). Thus, mutations potentially play a role in the formation of new MAT alleles under natural conditions. In the present study, the *HD1* gene mutations were found in *Pst* populations of Shaanxi, Henan, Jiangsu, Xinjiang and Sichuan provinces, especially Sichuan; the *HD2* gene mutations were observed in the *Pst* populations in Tibet and Sichuan (Table 6). It is hypothesized that *Pst* in certain areas in which annual infection cycle can complete and *Pst* mutations could be prone to occur (e.g. Sichuan), new *HD* gene types possibly formed through the accumulation of mutations.

**Table 6.** The mutation rate of mutational *HD1* and *HD2* gene sequences (testing sequences) in *Puccinia striiformis* f. sp. *tritici* isolates collected in China.

| Isolate code | Province | *HD* gene type | Identity (%) | Mutation rate (%) |
| --- | --- | --- | --- | --- |
| 1 | Tibet | *PstHD2-w2* | 99.88 | 0.12 |
| 2 |  | *PstHD2-w2* | 99.88 | 0.12 |
| 3 | Shaanxi | *PstHD1-e1* | 99.66 | 0.34 |
| 4 | Henan | *PstHD1-e5* | 98.70 | 1.30 |
| 5 |  | *PstHD1-e8* | 99.86 | 0.14 |
| 6 | Jiangsu | *PstHD1-e8* | 99.86 | 0.14 |
| 7 | Xinjiang | *PstHD1-e8* | 99.86 | 0.14 |
| 8 | Sichuan | *PstHD1-e8* | 99.58 | 0.42 |
| 9 |  | *PstHD1-e8* | 99.31 | 0.69 |
| 10 |  | *PstHD1-e8* | 99.58 | 0.42 |
| 11 |  | *PstHD1-e8* | 97.22 | 2.78 |
| 12 |  | *PstHD1-e8* | 99.03 | 0.97 |
| 13 |  | *PstHD1-e8* | 99.58 | 0.42 |
| 14 |  | *PstHD2-w3* | 99.87 | 0.13 |
| 15 |  | *PstHD2-w3* | 99.74 | 0.26 |
| 16 |  | *PstHD2-w3* | 99.60 | 0.40 |

The gene *PshHD1*-*e1*, *PshHD1-e2*, *PshHD2-w1* and *PshHD2-w2*, developed from the genome of *Psh*, are specific *HD* genes for *Psh*. The formation of this gene may be resulted from the directional host selection pressure to the pathogen. However, the identity between gene *PshHD1*-*e1* and the *PstHD1*-*e9* of *PstHD1* genes reached up to 81.63%, which was higher than that between some *Pst HD1* gene sequences (75.68% for *PstHD1*-*e1* and *PstHD1*-*e8*). The same is true for the *HD2* gene. The identity between *PshHD2-w2* and *PstHD2-w4* can reach up to 79.42%, which is higher than that between some *HD2* gene sequences of *Pst* (75.67% for *PstHD2-w8* and *PstHD2-w10*). *Puccinia striiformis* 11-281 containing the gene *PstHD1-e2* used grass as a host (Table 7). Accumulating evidence indicates that various races (or variants) are generated by genetic recombination or somatic recombination between races, and between *Pst* and *formae speicales* (f. spp.) of *P. striiformis* on grasses [47–51]. Therefore, it is speculated that in the natural *Pst* population, some of the *HD* types of *Pst* may be derived from other formae speciales of *P. striiformis*, and that, in turn, the *HD* genes of other formae speciales of *P. striiformis* are possibly introduced into *Pst* through genetic recombination, resulting in many of *HD* gene types with large sequence differences in *Pst*.

**Table 7.** Distribution of *Puccinia striiformis HD* genes genomes released worldwide.

| **HD gene** |  | **Origin of isolate<sup>a</sup>** | **Host** |
| --- | --- | --- | --- |
| *PstHD1-e1* | *PstHD2-w1* | China, USA, Canada | wheat |
| *PstHD1-e2* | *PstHD2-w2* | China, India, USA, Canada, Germany, Poland, New Zealand, Ethiopia, Zimbabwe, Kenya, South Africa | wheat, Grass, Triticale |
| *PstHD1-e3* | *PstHD2-w3* | China, India, USA, Canada, Germany, Poland Ethiopia, Zimbabwe, Kenya, Ethiopia, South Africa, | wheat |
| *PstHD1-e4* | *PstHD2-w4* | China, India, Turkey, USA, Canada, Australia, Chile, Denmark, UK, France, Germany, Croatia, Czechia, Austria, Poland, Bulgaria, Italy, Slovakia, Belgium, Romania, Serbia | wheat |
| *PstHD1-e5* | *PstHD2-w5* | China, USA, Canada, Australia, New Zealand, Chile, UK, France, Sweden, Poland | wheat |
| *PstHD1-e6* | *PstHD2-w6* | China | wheat |
| *PstHD1-e7* | *PstHD2-w7* | China, Pakistan, Ethiopia, | wheat |
| *PstHD1-e8* | *PstHD2-w8* | China, India, Turkey, Eritrea, Denmark, UK, Germany, Croatia, France, Austria, Poland, Czechia, Bulgaria, Italy, Slovakia, Belgium, Romania, Serbia, Sweden, | wheat |
| *PstHD1-e9* | *PstHD2-w9* | China | wheat |
| *PstHD1-e10* | *PstHD2-w10* | China, India | wheat |
| *PstHD1-e11* | *PstHD2-w11* | China, Pakistan, USA, Poland, Sweden, Ethiopia, Eritrea | wheat |

<sup>a</sup> The date referred to this study, Holden et al. [52], and NCBI Database.

In the present study, we demonstrated that the mating system of *Pst* is a tetrapolar mating system. Notably, most studies suggest that the mating system in the Basidiomycota may produce multiple mating type specificities in one or two unlinked mating type loci to improve outbreeding potential, so to allow the diversity of isolates in the population to be adequately maintained in the process of evolution [36, 53]. Thus, we suggest that the formation of diverse *HD* gene types in natural *Pst* populations may be an effective gene fragment resource for the evolution of multiple alleles in the locus, resulting in efficient outbreeding reproduction.

In this study, it is confirmed that the *HD* gene plays a biological function during sexual reproduction of *Pst* (Fig 2). Moreover, there were other MAT genes that are unlinked to *HD* locus and involved in sexual reproduction other than *HD* genes in *Pst* (Fig 2 and Table 2). A study by Cuomo *et al.* [54] revealed the presence of pheromone precursor and pheromone receptor genes in the genome of *Pst* by sequence alignment, such as genes *PstSTE3.1*, *PstSTE3.2*, *PstSTE3*.*3*, *Pstmfa2*, *Pstmfa*. Therefore, it is speculated that other *MAT* genes involved in sexual reproduction of *Pst* are possibly pheromone and pheromone receptor genes.

### Sexual reproduction responsible for high genetic diversity of natural *Pst* populations

Sexual reproduction is an important way to complete assortment of *HD1* (*HD2*) genes by selfing and generate new *HD1* (*HD2*) gene combination types by crossing in *Pst* (Figs 7A and 7B). In selfing system, each of a two-celled, diploid teliospore (2n) germinate four basidiospores. Two contain the same *HD* gene type and other two carry another identical *HD* gene type distinguishing from the former. The mating of two identical *HD* gene types, such as *PstHD-A* or *PstHD-B*, is not functional for fertilization, resulting in sterile. However, the mating occurs different *HD* gene types, such as *PstHD-A* and *PstHD-B*, nearly one half of which are fertile and can produce aecia, and the other are sterile and are not able to generate aecia (Fig 7A). In crossing system, the mating of two *HD* genes that are derived from different races, such as *PstHD-A* and *PstHD-D*, or *PstHD-B* and *PstHD-C*, can produce new *HD* gene recombination in an aeciospore (n+n) of progenies (Fig 7B). However, whether *HD* gene recombination, such as *PstHD-A* and *PstHD-C*, or *PstHD-B* and *PstHD-D* could occur by the mating, which is not undetermined. In addition, in this study, only one kind of *HD1* gene was detected in each of all nectar (pycniospores, n) samples of single pycnium, which indicated that *HD1* alleles were separated during the pycniospore stage of *Pst*. Whether the pair of *HD1* alleles of *Pst* is located in two nuclei at aeciospore stage, and it remains to be further investigated.

**Figure 7.**
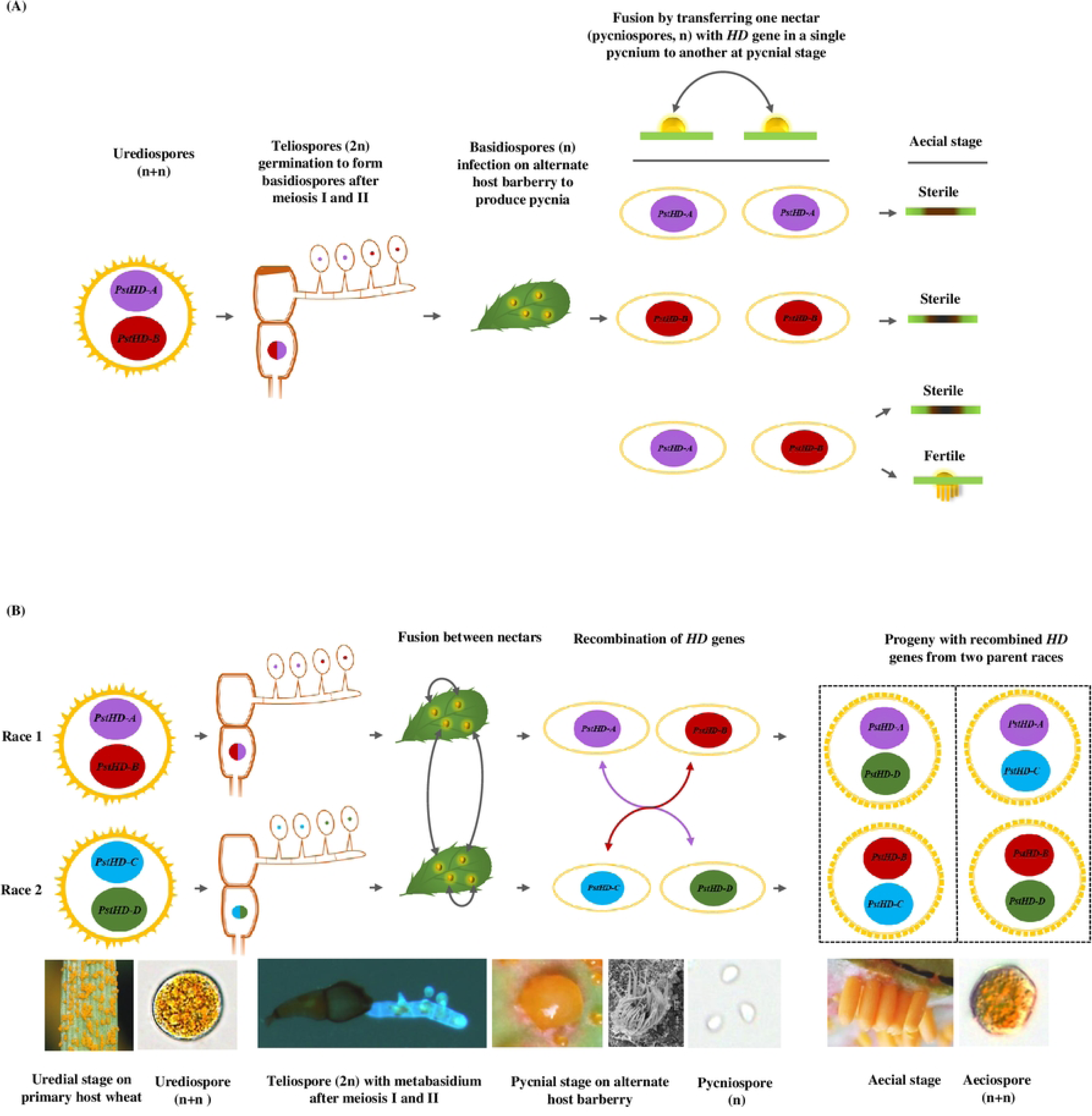
A skeleton illustrating tetrapolar system of *Puccinia striiformis* f. sp. *tritici* sexual reproduction. (a) Selfing. (b) Crossing.

There are abundant *HD1* (*HD2*) gene combination types in Chinese *Pst* population. In this study, we detected at least 30 types in *Pst* populations from eleven provinces of China (S6 Table). Combinations of one *HD1* (*HD2*) gene with multiple different of *HD1* (*HD2*) gene types were observed in common in overall population, which was exemplified by *PstHD1*-*e1* that was detected to combine with at least seven types of *HD1* genes (S7 Table). Thus, it is deduced that *HD1* (*HD2*) gene recombination could occur between different isolates in natural *Pst* populations in China.

Over the past one decade, we have investigated that many Chinese barberry *(Berberis* spp.) species were identified as alternate hosts for *Pst* and widely distribute in many provinces of China [23, 55–58]. Importantly, it has been demonstrated that under natural conditions *Pst* can infect susceptible barberry to complete sexual cycle in many Chinese provinces, including Gansu, Shaanxi, Qinghai, and Tibet [23–27], and some of which susceptible barberry provide aeciospore inoculum to cause stripe rust on wheat [25, 27]. Those studies showed that a potential sexual outcrossing between isolates could occur to give rise to the abundance of *HD1* (*HD2*) gene combination types, resulting in the increase of genetic diversity in natural *Pst* populations. Under laboratory conditions, it has been testified that either of selfing and sexual outcrossing of *Pst* can lead to genetic recombination and produce diverse progenies with different virulence from the parental isolates (Table 4) [59–62]. Furthermore, we observed there were virulence differences among the most isolates with same or different *HD1* (*HD2*) gene combination types in the natural *Pst* population. Potential approaches for leading to virulence variation are needed to prove.

The emergence of novel *Pst* races that overcome the resistance of wheat cultivars is a significant contributing factor to the occurrence of large-scale epidemics [12, 14]. It is crucial that we place a strong emphasis on the new *Pst* races produced through sexual recombination, as this can enable us to develop appropriate strategies for the control of stripe rust. These strategies can be proposed in advance to mitigate the risk of disease epidemics.

### Population structure and genetic diversity of *Pst* populations through MAT gene *HD*

In this study, structure and cluster analyses of the *HD1* gene of overall *Pst* population indicated that the *Pst* populations in Tibet and Xinjiang were considered as respectively isolated from those of other provinces, which was consistent with previous studies [7, 21, 63–64]. But compared with Tibet, Xinjiang had relatively more frequent exchanges with other regions of China. The isolation of the Tibetan population from other provincial populations can be attributed to the formidable Himalayan mountains, which serve as a barrier, restricting the geneflow between Tibet and the other provinces. *HD1* genes (*PstHD1*-*e2*, *PstHD1*-*e4*) and its gene combination T1 (*PstHD1*-*e2* + *PstHD1*-*e4*) are unique duet to detection only in Tibetan population but not in other provincial populations of China. Through data analysis, we found that the genes *PstHD1-e2* and *PstHD1-e4* were distributed in the Asia, America, Europe, Africa and Oceania (Table 7) [52]. This result indicated that the *Pst* population in Tibet possibly has gene drift with *Pst* population worldwide. Our study provides novel evidence that supports the hypothesis that the Himalayan region is the probable centre of origin of *Pst* [16].

Xinjiang population is isolated from other provincial population based on the detection of the *HD1* gene combination type T4 (*PstHD1*-*e3*+*PstHD1*-*e8*) only in Xinjiang but not in other provincial populations, which is accordance with early conclusion as reported by Li and Zeng [7]. Nevertheless, either of two *HD1* genes *PstHD1*-*e3* and *PstHD1*-*e8* distributes not only in Xinjiang population but also in other provincial populations in China, which was inferred that there could be gene drift among populations of Xinjiang and other provinces in China. This inferring is in consistent with a study by Wan *et al.* [65] reported that Xinjiang population may not be completely isolated due to the existence of weak genetic exchange between Xinjiang and Gansu. The genes *PstHD1-e3* and *PstHD1-e8* were widely distributed in Asia, America, Europe and Africa through data analysis (Table 7) [52]. According to this study, it was found that *PstHD1-e8* is predominant among HD1 gene type in the *Pst* population in China. Because *PstHD1-e8* was the most widespread (detected in 10 provincial populations), with the highest percentage (30%) and the largest number of combination types (combined with at least five *HD1* genes). Based on the above results, it was inferred that Xinjiang may be the main channel of communication between China and groups in the world. Recent study conducted by Awais et al [66], showed the evidence of *Pst* invasion from Central Asia to Xinjiang, reflecting new races may produce in this region due to crossing among local and Central Asian genotypes.

Population genetic relationship of *Pst* based on the clustering of *HD1* gene of overall *Pst* population was generally consistent with those of molecular data reported previously [8, 21, 67–70]. Thus, *HD1* gene is essential for the process of sexual recombination, and conversely, molecular markers are independent to this process due to the nature of neutrality. Whereas, both *HD1* gene and molecular markers showed similar genetic relationship for Chinese *Pst* natural population. *HD* genes were useful molecular markers for the analysis for genetic diversity of the stripe rust pathogen. In the present study, many *HD1* gene sequences were still unknown in natural *Pst* population, and thus more accurate and precise analyses are needed.

Mutations of *HD1* gene were mostly detected in Sichuan *Pst* population, rather than Gansu or other provinces of key variable regions, which are unexpected. Historically, although some of new races have been first detected in this province [71–72], which was minor. Most recently, molecular data analyses revealed that Sichuan population exhibited the highest genetic diversity compared to other Chinese provincial populations, with a strong signal of genetic recombination [21]. However, although alternate host barberry native to Sichuan have been identified [23], no evidence has been reported the existence of sexual reproduction of the rust in this region. Moreover, whether mutations of *HD1* gene is related to local climate and surroundings environment conditions, which is needed to confirm.

Chinese *Pst* population possesses a significantly high genetic diversity based on different molecular marker methods [15, 18, 21, 73], compared with clonal populations from Europe [14,73], America [20, 75], and Australia [76–77]. High genetic diversity of Chinese *Pst* population can be exemplified by a *Pst* population from Tianshui of Gansu in China in one year that was seven folds greatly higher than those collected in 20 years in France in the richness of genetic diversity [18]. Therefore, in the case of the unknown of sexual stage of the rust it has been hypothesized that Chinese *Pst* population could be produced sexually [18]. Whereas, with the finding of barberry (*Berberis*) being identified as alternate hosts for *Pst* [22], under natural conditions in China it has been demonstrated that *Pst* infects wild barberry species to complete sexual cycle [23–24, 26, 78], and that susceptible barberry species play a role in providing aeciospores inoculum to trigger stripe rust on wheat [25, 27], but not in American and European countries [79–80]. Additionally, sexual reproduction of *Pst* can give rise to a high level of pathogenic variation and is considered as an important approach of new variants of the rust [59, 62]. In this study, regardless of *HD* gene types or *HD* gene combination types Gansu, as a core oversummering variable region, *Pst* population of which was maximum among overall provincial populations, followed by those of Qinghai and/or Shaanxi. Thereby, high level of the abundance of gene types and recombination types of *HD* gene in Chinese *Pst* population could be attributed from the frequent occurrence of sexual cycles due to genetic recombination at sexual stage.

## Methods

### Plant materials

Susceptible barberry seeds (*Berberis shensiana*) were collected from Baoji, Shaanxi, China. The seeds were dried at room temperature in a laboratory ensure to prevent them direct sunlight exposure. After drying, the seeds were stored in a desiccator containing silica gels in a freezer at a temperature of 4°C until their intended use. Barberry seedlings were cultivated using the method described by Zhao et al. [23]. Inoculation was conducted when the barberry seedlings reached the four to six-leaf stage. Wheat cultivar Mingxian 169, a highly susceptible for all known Chinese races of *Pst*, were used for establishing a pure isolate from a stripe rust sample and used it for spore multiplication. A set of differential hosts, including 25 wheat lines carrying single *Yr* gens (S8 Table), was used for analyzing race identification and virulence variation. Wheat cultivar Mingxian 169 was used a positive control in all tests. Seven to ten wheat seeds were grown in a small plastic pot filled with commercial potting mix (Inner Mongolia Mengfei Biotech Co., Ltd., Huhhot, Inner Mongolia), and cultivated at 20-25℃ in a condition-controlled rust-free growth chamber. A dual photoperiod regime of 16 h light and 8 h darkness was used. Eight-day-od wheat seedlings were used for inoculation and increasing spores.

### Fungal isolates

In this study, we utilized stored spores of races CYR23 and CYR32 from our laboratory. Additionally, stripe rust leaf samples of wheat were collected from wheat plants in eleven provinces, including Gansu, Guizhou, Henan, Hubei, Jiangsu, Qinghai, Shaanxi, Sichuan, Xinjiang, Yunnan, and Tibet. Similarly, barley leaf samples were collected from Tibet (S2 Table). For obtaining a pure isolate, an individual leaf sample was put onto 2-layer wetted filter papers inside a plate and kept at room temperatures for 1 h for hydration. After hydration, a single uredinium (sorus) was picked from a leaf specimen to inoculate onto a leaf of seedlings of wheat cv. Mingxian 169 for obtaining a pure isolate. Likewise, using the same method as described above, obtainment of a pure culture of *P. striiformis* f. sp. *hordei* (*Psh*, the causal pathogen of barley stripe rust) was conducted by inoculating a leaf of barley cv. Guoluo seedlings, highly susceptible to all known Chinese *Psh* races, with an individual uredinium of a barley stripe rust leaf sample. Inoculated plants in a pot were isolated by covering a transparent plastic cylinder (binding film, Deli Group, Zhejiang, China). After sprayed with deionized water, inoculated plants were put in a tray and incubated for 24 h at 10℃ with 100% relative humidity in the dark in a dew chamber (I-36DL; Percival Scientific, Inc., IA, USA). After incubation, plants were moved to a condition-controlled growth chamber with a dual photoperiod regime of 16 h light at 16℃ and 8 h darkness at 13℃ for sporulation. Fresh urediniospores were collected and kept inside a dessicator with silica gels at 4℃ in a refrigerator until use.

### Preparation and teliospore formation of single-urediniospore isolate

Single-urediniospore isolate of *Pst* races, CYR23 and CYR32, was established by picking a single urediniospore under a dissecting microscope according to the method described by Zhang *et al.* [80]. Spores were increased by re-inoculating wheat cv. Mingxian 169. To obtain teliospores, the sporulating wheat plants were continuously cultivated to induce teliospore formation by alternating 16℃ and 25℃ temperatures. When telia obviously appeared, leaves were collected from wheat plants, dried completely, and then kept in a silica gels-filling dessicator in low temperatures (4-5℃) in a refrigerator for later use.

### Genomic DNA extraction, RNA extraction and cDNA synthesis

Genomic DNA was extracted from urediniospores (∼20 mg) according to the CTAB method [62, 81]. Total RNA was extracted from wheat (cv. Mingxian 169) leaves infected with each of *Pst* isolates, and barley (cv. Guoluo) infected leaves with each of *Psh* isolates at 10 days post-inoculation (dpi) using a Quick RNA isolation Kit (Huayueyang Biotechnology (Beijing) Co., Ltd., China) according to the manufacturer’s instructions. First-strand cDNA was synthesized using a RevertAid^TM^ First Strand cDNA Synthesis Kit (Thermo Fisher Scientific (China) Co., Ltd., Shanghai). Quality and quantity of genomic DNA and cDNA were measured using a spectrophotometer (NanoDrop 2000, Thermo Fisher Scientific, MA, USA). The diluted DNA and cDNA solutions were stored at -20℃ until use.

### Cloning, physical arrangement and specific primer design for *HD* genes

To design sequences of *HD* gene primers and understand conserved characteristics of *Pst* and *Psh*, multi-sequence alignments of genome sequences of 17 *Pst* and *Psh* isolates (S9 Table), downloaded from NCBI (https://www.ncbi.nlm.nih.gov/), were used to obtain sequences of *HD* genes of *Pst* and *Psh* (S10 Table). Based on conserved sequences at two ends of *HD1* and *HD2* genes of *Pst* and *Psh*, *HD* primers were designed using Primer Premier 5.0 (Premier Biosoft, Canada). The designed primers were used to amplify the target genes of *HD1* and *HD2* in the genomes of 29 *Pst* isolates and 3 *Psh* isolates using PCR amplification in a thermal cycler (VeritiPro, Applied Biosystems Inc., CA, USA). PCR program was run by pre-denaturing for 2 min at 94℃; 30 cycles consisting of denaturing for 30 s at 94℃, annealing for 30 s at 55℃, and extension for 75 s at 72℃; and a final extension for 2 min at 72℃ prior to ending at 4℃. The amplicons were purified with an Original TA Cloning^TM^ Kit (Thermo Fisher Scientific, MA, USA) and sequenced by Sangon Biotech (Shanghai) Co., Ltd.

To understand whether *HD1* and *HD2* genes are linked, according to conserved sequences of the middle regions of *HD1* and *HD2* genes, primer 1/ primer 4 pair, or primer2/primer3 pair in this region was designed (S2 Fig and S11, S12 Tables), and used to amplify using the target amplicons (mentioned above) as DNA templates, respectively. In the light of sequences of cloned *HD1* and *HD2* genes of *Pst* and *Psh*, *HD1* and *HD2* gene-specific primers were designed respectively, and synthesized by Sangon Biotech (Shanghai) Co., Ltd. PCR amplicons were obtained by running the same program as above, and sequenced, sequences of which were compared, and analyzed with those corresponding to *HD1* gene and *HD2* gene.

### *HD* gene detection based on PCR amplification

The primers specific for *HD1* and *HD2* genes were used for PCR amplification (S10 Table), which was carried out in a total volume of 20 µl consisting of 10 µl of 2×Es *Taq* Master Mix (Dye), 2.0 µl of template DNA (50 ng/µl), 1µl of each of forward and reverse primers (10 µM), and 6 µl of double distilled water (ddH_2_O). The amplification conditions comprised of a pre-denaturation at 94°C for 2 min, followed by 30 cycles consisting of denaturation at 94°C for 30 s, annealing at 55°C for 30 s, and extension at 72°C for 45 s, and a final extension at 72°C for 2 min. The PCR products were separated by 1% (m/vol.) agarose gel electrophoresis, and photographed using a gel imaging system (GelDoc XR, Bio-Rad, CA, USA). Three repetitions were accomplished for ensuring the consistent of PCR amplicons. The PCR amplicons were sequenced using an ABI sequencer (3730xl DNA Analyzer, Life Technologies Inc., Shanghai, China).

### Barberry inoculation

Inoculation of excised barberry young leaves with basidiospores, produced from germinated teliospores, was performed to initiate pycnial stage, using method as described by Zheng *et al.* [48]. Prior to inoculation, leaf segments bearing teliospores were soaked with deionized water for completely hydration and put onto 2% (m/vol.) water agar plate on the top cover of a Petri dish at 10℃ until initial germination of teliospores started. The top cover was turned downward to detached barberry leaves that each were wrapped around a petiole of a detached leaf with water-saturated cotton patches on the bottom of the dishes with a layer of plastic gauze (S6A Fig). The Petri dishes were sealed using parafilm after spraying, and kept in an incubator with an alternate photoperiod system of 16 h light and 8 h darkness at 10°C for 3 d. After incubation, the dishes were maintained in the incubator with dual photoperiod regimes of 16 h light time at 16℃ and 8 h darkness at 13℃ until the emergence of pycnial signs (S6B Fig). During cultivation of inoculated leaves, timely adding water to cotton patches was implemented for maintaining detached leaves alive.

### Identification of mating system and *HD* genes exchange

To determine that mating system, and the *HD* genes exchange of *Pst* during the process of sexual reproduction, detached young barberry leaves were inoculated with basidiospore. When an obvious nectar (pycniospores) was observed on an ostiole of a pycnium after inoculation *in vitro* as described above, a 2-μl of nectar produced in a pycnium was pipetted into a microcentrifuge tube for PCR amplification with *Pst HD* specific primers for detecting *HD* gene in the pycnium. The transfer of nectar from one pycnium to another was conducted to ascertain whether nectars can fuse for fertilization, thereby completing sexual reproduction. Thus, nectar transfers were designed for confirming mating system by fusion between nectars produced from pycnia carrying same and different *Pst* MAT genes. Two sexual populations were constructed by selfing and hybridization to determine exchange of MAT genes. One was obtained from selfing race CYR32 on barberry leaves, and the other was constructed by crossing race CYR23 and race CYR32 on the barberry leaves, respectively.

All mated nectars on pycnial lesions were labelled, and the barberry leaves were maintained inside the plate, and periodically observed. When aecia initially emerged on the backside of the leaves, to avoid aeciospores being ineffective, the leaves were then overturn upward for aecia growing longer in length and the lid was covered the half of the plate. All mated pycnia were recorded with/without aecial production. Approximately one third of an individual aecium (aecial cup) was excised, put onto a clean glass slide, crushed gently to expose aeciospores. Aeciospores were then added two drops (∼20 μl) of deionized water to make spore suspension. The suspension was inoculated leaves of a seedling plant of wheat cv. Mingxian 169 using the same routine method as described above. The inoculated wheat plants were cultivated until urediniospores emergence in a rust-free growth chamber under same temperatures and photoperiod conditions as above. The remaining (∼2/3) of an aecium (aecial cup) were collected, and aeciospores of which were used for detecting *HD* genes using *HD* specific primers. The same method of genomic DNA extraction and identical PCR amplification system, as mentioned above, were used.

### Virulence phenotyping

The virulence phenotypes (VPs) of isolates were determined using a set of 25 single-*Yr* wheat lines (S8 Table). Seedlings of wheat lines were inoculated with urediospores according to the method described by Zhao *et al.* [23]. After inoculation, the plants are transferred to a growth chamber with the same conditions as above. The 0-to-9 scale described by Line and Qayoum [83] was used to record infection types (ITs) 18 to 20 d after inoculation. Virulence test was repeated twice. ITs 0-to-6 were considered as avirulent (A), while 7-to-9 as virulent (V).

### Detection of MAT genes in natural population

To understand *HD* genes of *Pst* and *Psh* natural population, and association with *Pst* population genetic diversity, all of *HD* gene specific primers, developed in this study, were used to detect *HD* genes of overall isolates of both rust pathogens from 11 provinces of China by PCR method. Same PCR amplification system and program were referenced as above.

### Data analysis

To evaluate virulence variation of progenies produced by selfing and hybridization of *Pst* races and natural *Pst* races, the VAT software was used to compare avirulence or virulence of each of progenies on each resistance locus of the 25 *Yr*-single gene lines with those of parental races.

To determine probability of successful fertilization via heterothallic nectar fusion, using the interval estimation of 0-1 distribution parameters, the confidence level with the probability of producing aecia of less than 100% was calculated when *α* is equal to 0.001 (*α*=0.001).

To evaluate the level of genetic diversity of natural population of *Pst* based on *HD* genes, the following analyses were performed. (1) STRUCTURE analysis: STRUCTURE 2.3.4 software [84] was analyzed based on the likelihood of 100,000 by setting the operating parameter *K* (from 2 to 11), and repeated 10 times. The best *K* value was determined by online tool Structure Harvester (https://taylor0.biology.ucla.edu/structureHarvester). Repeated sampling analysis was accomplished using CLUMPP1.1.2 [84], and DISTRUCT software was used to display graphically the results. (2) Discriminant analyses of principal components (DAPC): In the R language environment, the ADEGENET program package was used to perform DAPC analysis based on multivariate and model-based Bayesian clustering methods [86]. (3) Cluster analysis: Population genetic analysis was carried out in POPGENE VERSION 1.31 to calculate summary statistics for multiple populations [86]. The genetic identity of multiple populations was estimated [87], and then a dendrogram was established using NTSYSpc software (ver. 2.10e).

## Supporting information

**S1 Fig. A drawing illustrating linkage between *HD1* gene and *HD2* gene in *Puccinia striiformis* f. sp*. tritici.*** (A) and *P. striiformis* f. sp. *hordei* (B).

**S2 Fig. The physical arrangement of *HD1-HD2* gene pair in *Puccinia striiformis* f. sp. *tritici* (*Pst*) and *P. striiformis* f. sp. *hordei* (*Psh*). (A) A skeleton illustrating physical arrangement of *HD1*-*HD2* gene pairs. (B)** Stained-agarose gels showing PCR amplified products of both *HD1*-*HD2* gene pairs in the genome of 11 *Pst* isolates and 2 *Psh* isolates with primer 1 and primer 4 for the *HD1* gene (I), and primer 2 and primer 3 for the *HD2* gene (II), respectively. The primers listed in S11 Table and S12 Table.

**S3 Fig. Homeodomain protein sequences in HD1 and HD2 transcription factors from mating type loci of *Puccinia striiformis* f. sp. *tritici*, *P. striiformis* f. sp. *hordei* and selected fungi of basidiomycetes.** (A-B). Homeodomain protein sequences in HD1 and HD2 transcription factors from *Puccinia striiformis* f. sp. *tritici* (*PstHD1-e1* to *PstHD1-e11*, *PstHD2-w1* to *PstHD2-w11*) and *P. striiformis* f. sp. *hordei* (*PshHD1-e1* and *PshHD1-e2*, *PshHD2-w1* and *PshHD2-w2*). (B-C) Homeodomain protein sequences in HD1 and HD2 transcription factors from *Puccinia striiformis* f. sp. *tritici*, *P. striiformis* f. sp. *hordei* and selected fungi of basidiomycetes listed in S1 Table. The red dotted line rectangle blocks indicated the three helical regions of the homeodomain proteins (I, II, and III) of HD1 and HD2 transcription factors, and the positions of the conserved amino acids of the DNA binding motif indicated by bold white font uppercases, respectively.

**S4 Fig. Stained-agarose gels showing composition of *HD1* genes of parental races (CYR 23 and CYR32) and progenies of *Puccinia striiformis* f. sp. *tritici* between a cross of both races based on detection of PCR amplification products with specific primers for four *PstHD1* genes (*PstHD1-e1*, *PstHD1-e3*, *PstHD1-e8*, and *PstHD1-e10*) (A), and four *PstHD2* genes (*PstHD2-w1*, *PstHD2-w3*, *PstHD2-w8*, and *PstHD2-w10*) (B), respectively.** M, DL2000 DNA Marker (Shanghai Yuanye Bio-Technology Co., Ltd., China).

**S5 Fig. Scatter-plot of the discriminant analysis of principal components (DAPC) on *HD1* genes of *Puccinia striiformis* f. sp. *tritici* isolates from 11 provinces of China.** The eigenvalues at the first two axes are represented at the bottom left corner. The genotype as dots and clusters as ellipses are represented in the graph at the right corner.

**S6 Fig. The culture of detached-leaves of barberry (*Berberis shensiana*) inoculated using basidiospores produced from germinated teliospores of *Puccinia striiformis* f. sp. *tritici* race CYR32 in the petri dish.** (A) Culture of detached-leaves of *Berberis* infected with basidiospores. (B) Close-up view of an obvious nectar (pycniospores) drop accumulating on an ostiole of a pycnium circled with a yellow dotted line in (A).

**S1 Table. HD1 and HD2 transcription factors of selected fungi of basidiomycetes.**

**S2 Table. Information of *Puccinia striiformis* f. sp. *tritici* (*Pst*) and *P. striiformis* f. sp. *hordei* (*Psh*) isolates collected in 11 provinces of China.**

**S3 Table. The *HD1* gene testing of *Puccinia striiformis* isolates collected from 11 provinces in China using specific primers of *HD1* gene.** ^a^ The testing used 13 pairs of *HD1* gene specific primers of *Puccinia striiformis* (S10 Table); E1=*PstHD1*-*e1*, E2=*PstHD1*-*e2*, E3=*PstHD1*-*e3*, E4=*PstHD1*-*e4*, E5=*PstHD1*-*e5*, E6=*PstHD1*-*e6*, E7=*PstHD1*-*e7*, E8=*PstHD1*-*e8*, E9=*PstHD1*-*e9*, E10=*PstHD1*-*e10*, E11=*PstHD1*-*e11*, E12=*PshHD1*-*e1*, E13=*PshHD1*-*e2*; 1=having hand, 0=no band; the highlighted indicates there are mutations in the sequence; no highlighting indicates there is no mutation in the sequence. ^b^ I, The samples of *Puccinia striiformis* f. sp. *hordei* (*Psh*) collected from barley. ^c^ II, The samples of *Puccinia striiformis* f. sp. *tritici* (*Pst*) collected from wheat.

**S4 Table. The *HD2* gene testing of *Puccinia striiformis* isolates collected from 11 provinces in China using specific primers of *HD2* gene.** ^a^ The testing used 13 pairs of *HD2* gene specific primers of *Puccinia striiformis* (S10 Table); W1=*PstHD2-w1*, W2=*PstHD2*-*w2*, W3=*PstHD2*-*w3*, W4=*PstHD2*-*w4*, W5=*PstHD2*-*w5*, W6=*PstHD2*-*w6*, W7=*PstHD2*-*w7*, W8=*PstHD2*-*w8*, W9=*PstHD2*-*w9*, W10=*PstHD2*-*w10*, W11=*PstHD2*-*w11*, W12=*PshHD2*-*w1*, W13=*PshHD2*-*w2*; 1=having hand, 0=no band; the highlighted indicates there are mutations in the sequence; no highlighting indicates there is no mutation in the sequence. ^b^ I, The samples of *Puccinia striiformis* f. sp. *hordei* (*Psh*) collected from barley. ^c^ II, The samples of *Puccinia striiformis* f. sp. *tritici* (*Pst*) collected from wheat.

**S5 Table. The mutation rate of mutational *HD1* and *HD2* gene sequences (testing sequences) in *Puccinia striiformis* f. sp. *tritici* isolates collected in China.**

**S6 Table. *HD* gene combination types of *Puccinia striiformis* f. sp. *tritici* in 11 provinces of China.**

**S7 Table. The number of *HD* gene combination types in natural population of *Puccinia striiformis* f. sp. *tritici* in China.**

**S8 Table. Wheat genotypes used in this study.**

**S9 Table. The information of *Puccinia striiformis* genome in data of the National Center for Biotechnology Information (NCBI) Database.**

**S10 Table. Primers used for cloning *HD* genes of *Puccinia striiformis* and gene testing.**

**S11 Table. Primers utilization for investigating the physical arrangement of the *HD1-HD2* gene pairs.** The primers listed in Table S12.

**S12 Table. Primers used for investigating physical arrangement of both *HD1*-*HD2* gene pairs.**

**S1 Data. The sequences of *HD1* gene and *HD2* gene in *Puccinia striiformis*.**

## Funding

This work was supported by National Key R&D Program of China (2021YFD1401000; 2024YFD1401000), National Natural Science Foundation of China (32272507, 32072358), and Natural Science Basic Research Plan in Shaanxi Province of China (2019JCW-18, 2020JCW-16).

## Author Contributions

JZ, SL, and ZK conceived and designed this study; ZD collected and identified samples; SL, MS, XC, and LY accomplished experiments; SL and JZ edited the manuscript.

## Competing interests

No declaration.

## Notes

### Competing Interest Statement

The authors have declared no competing interest.

## References

1. Beddow JM, Pardey PG, Chai Y, Hurley TM, Kriticos DJ, Braun HJ, Park RF, Cuddy WS, Yonow T. Research investment implications of shifts in the global geography of wheat stripe rust. Nature Plants. 2015; 1:15132. 10.1038/nplants.2015.132 PMID: 27251389.

2. Wang M, Wan A, Chen X. Race Characterization of *Puccinia striiformis* f. sp. *tritici* in the United States from 2013 to 2017. Plant Disease. 2022; 106(5):1462–1473. 10.1094/PDIS-11-21-2499-RE PMID: 35077227.

3. Roelfs AP, Singh RP, Saari EE. Rust Diseases of wheat: concepts and methods of disease management. CIMMYT: Mexico City, Mexico, 1992; pp. 1–81.

4. Chen XM. Epidemiology and control of stripe rust [*Puccinia striiformis* f. sp. *tritici*] on wheat. Canadian Journal of Plant Pathology. 2005; 27:314–37. 10.1080/07060660509507230.

5. Wellings C. Global status of stripe rust: A review of historical and current threats. Euphytica. 2011; 179:129–41. 10.1007/s10681-011-0360-y.

6. Fang CT. Physiologic specialization of *Puccinia glumarum* Erikss. and Henn. in China. Phytopathology. 1944; 34:2010–14.

7. Li ZQ, Zeng SM. Wheat Rusts in China. Beijing: China Agriculture Press. 2002.

8. Ma ZH. Researches and control of wheat stripe rust in China. Journal of Plant Protection. 2018; 45:1–6.

9. Liu WC, Wang BT, Zhao ZH, Li Y, Kang ZS. Historical review and countermeasures of wheat stripe rust epidemics in China. China Plant Protection. 2022; 42(6):21–7+41.

10. Wellings CR, Mcintosh RA. *Puccinia striiformis* f. sp. *tritici* in Australasia: pathogenic changes during the first 10 years. Plant Pathology. 1990; 39:316–25. 10.1111/j.1365-3059.1990.tb02509.x.

11. Chen XM, Penman L, Wan AM, Cheng P. Virulence races of *Puccinia striiformis* f. sp. *tritici* in 2006 and 2007 and development of wheat stripe rust and distributions, dynamics, and evolutionary relationship of races from 2000 to 2007 in the United States. Canadian Journal of Plant Pathology. 2010; 32:315–33. 10.1080/07060661.2010.499271.

12. Wan AM, Chen XM, He ZH. Wheat stripe rust in China. Australian Journal of Agricultural Research. 2007; 58:605–19. 10.1071/AR06142.

13. Han DJ, Wang QL, Chen XM, Zeng QD, Wu JH, Xue WB, Zhan GM, Huang LL, Kang ZS. Emerging *Yr26*-virulent races of *Puccinia striiformis* f. sp. *tritici* are threatening wheat production in the Sichuan Basin, China. Plant Disease. 2015; 99:754–60. 10.1094/PDIS-08-14-0865-RE PMID: 30699539.

14. Hovmøller MS, Walter S, Bayles RA, Hubbard A, Flath K, Sommerfeldt N, Leconte M, Czembor J, Rodriguez-Algaba J, Thach T, Hansen JG, Lassen P, Justesen AF, Ali S, de Vallavieille-Pope C. Replacement of the European wheat yellow rust population by new races from the centre of diversity in the near-Himalayan region. Plant Pathology. 2016; 65:402–11. 10.1111/ppa.12433.

15. Sharma-Poudyal D, Bai Q, Wan A, Wang M, See D, Chen X. Molecular characterization of international collections of the wheat stripe rust pathogen *Puccinia striiformis* f. sp. *tritici* reveals high diversity and intercontinental migration. Phytopathology. 2020; 110(4):933–42. 10.1094/PHYTO-09-19-0355-R PMID: 31895005.

16. Ali S, Gladieux P, Leconte M, Gautier A, Justesen AF, Hovmøller MS, Enjalbert J, de Vallavieille-Pope C. Origin, migration routes and worldwide population genetic structure of the wheat yellow rust pathogen *Puccinia striiformis* f. sp. *tritici*. PLoS Pathogens. 2014;10(1):e1003903. 10.1371/journal.ppat.1003903 PMID: 24465211.

17. Mboup M, Leconte M, Gautier A, Wan AM, Chen W, de Vallavieille-Pope C, Enjalbert J. Evidence of genetic recombination in wheat yellow rust populations of a Chinese oversummering area. Fungal Genetics and Biology. 2009; 46(4):299–307. 10.1016/j.fgb.2008.12.007 PMID: 19570502.

18. Duan X, Tiller A, Wan A, Leconte M, de Vallavieille-Pope C, Enjalbert J. *Puccinia striiformis* f. sp. *tritici* presents high diversity and recombination in the over-summering zone of Gansu, China. Mycologia. 2010; 102:44–53. 10.3852/08-098 PMID: 20120228.

19. Cheng P, Chen X M. Virulence and molecular analyses support asexual reproduction of *Puccinia striiformis* f. sp. *tritici* in the U.S. Pacific Northwest. Phytopathology. 2014; 104:1208–20. 10.1094/PHYTO-11-13-0314-R PMID: 24779354.

20. Holtz MD, Kumar K, Zatinge JL, Xi K. Genetic diversity of *Puccinia striiformis* from cereals in Alberta, Canada. Plant Pathology. 2014; 63:415–24. 10.1111/ppa.12094.

21. Awais M, Ali S, Ju M, Liu W, Zhang GS, Zhang ZD, Li ZJ, Ma XY, Wang L, Du ZM, Tian XX, Zeng QD, Kang ZS, Zhao J. Countrywide inter-epidemic region migration pattern suggests the role of southwestern population to wheat stripe rust epidemics in China. Environmental Microbiology. 2022; 24:4684–701. 10.1111/1462-2920.16096 PMID: 35859329.

22. Jin Y, Szabo LJ, Carson M. Century-old mystery of *Puccinia striiformis* life history solved with the identification of *Berberis* as an alternate host. Phytopathology. 2010; 100:432–5. 10.1094/PHYTO-100-5-0432 PMID: 20373963.

23. Zhao J, Wang L, Wang ZY, Chen XM, Zhang HC, Yao JN, Zhan GM, Chen W, Huang LL, Kang ZS. Identification of eighteen *Berberis* species as alternate hosts of *Puccinia striiformis* f. sp. *tritici* and virulence variation in the pathogen isolates from natural infection of barberry plants in China. Phytopathology. 2013; 103:935–40. 10.1094/PHYTO-09-12-0249-R PMID: 23514262.

24. Wang MN, Wan AM, Chen XM. Barberry as alternate host is important for *Puccinia graminis* f. sp. *tritici* but not for *Puccinia striiformis* f. sp. *tritici* in the U.S. Pacific Northwest. Plant Disease. 2015; 99:1507–16. 10.1094/PDIS-12-14-1279-RE PMID: 30695965.

25. Chen W, Zhang ZD, Ma XY, Zhang GS, Yao Q, Kang ZS, Zhao J. Phenotyping and genotyping analyses reveal the spread of *Puccinia striiformis* f. sp. *tritici* aeciospores from susceptible barberry to wheat in Qinghai of China. Frontiers in Plant Science. 2021; 12:764304. 10.3389/fpls.2021.764304 PMID: 34975948.

26. Liu Y, Chen XY, Ma Y, Meng ZY, Wang FL, Yang XJ, Chen XF, Li XM, Kang ZS, Zhao J. Evidence of roles of susceptible barberry in providing (primary) inocula to trigger stripe rust infection on wheat in Longnan, Gansu. Acta Phytopathologica Sinica. 2021; 51:366–80. 10.13926/j.cnki.apps.000704.

27. Zhao YY, Huang XL, Li Q, Huang LL, Kang ZS, Zhao J. Virulence phenotyping and molecular genotyping reveal high diversity within and strong gene flow between the *Puccinia striiformis* f. sp. *tritici* populations collected from barberry and wheat in Shaanxi Province of China. Plant Disease. 2023; 107:701–12. 10.1094/PDIS-12-21-2713-RE PMID: 35869588.

28. Beukeboom L, Perrin N. The Evolution of Sex Determination. Oxford UK: Oxford University Press. 2014.

29. Nieuwenhuis BP, James TY. The frequency of sex in fungi. Philosophical Transactions of The Royal Society B-biological Sciences. 2016; 371(1706):20150540. 10.1098/rstb.2015.0540 PMID: 27619703.

30. Heitman J, Sun S, James TY. Evolution of fungal sexual reproduction. Mycologia. 2013; 105:1–27. 10.3852/12-253 PMID: 23099518.

31. Ene I V, Bennett R J. The cryptic sexual strategies of human fungal pathogens. Nature Reviews Microbiology. 2014; 12(4):239–51. 10.1038/nrmicro3236 PMID: 24625892.

32. Wilson AM, Wilken PM, Wingfield MJ, Wingfield BD. Genetic networks that govern sexual reproduction in the *Pezizomycotina*. Microbiology and Molecular Biology Reviews. 2021; 85(4):e0002021. 10.1128/MMBR.00020-21 PMID: 34585983.

33. Lee S C, Ni M, Li W J, Shertz C, Heitman J. The evolution of sex: a perspective from the fungal kingdom. Microbiology and Molecular Biology Reviews. 2010; 74(2):298–340. 10.1128/MMBR.00005-10 PMID: 20508251

34. Sun S, Coelho MA, David-Palma M, Priest SJ, Heitman J. The evolution of sexual reproduction and the mating-type locus: links to pathogenesis of *Cryptococcus* human pathogenic fungi. Annual Review of Genetics. 2019; 53:417–44. 10.1146/annurev-genet-120116-024755 PMID: 31537103.

35. Kück U, Pöggeler S. Gryptic sex in fungi. Fungal Biology Reviews. 2009; 23:86–90. 10.1016/j.fbr.2009.10.004.

36. Kües U, James TY, Heitman J. 6 Mating Type in Basidiomycetes: Unipolar, Bipolar, and Tetrapolar Patterns of Sexuality. Springer Berlin Heidelberg. 2011. 10.1007/978-3-642-19974-5_6.

37. Coelho MA, Bakkeren G, Sun S, Hood ME, Giraud T. Fungal Sex: The Basidiomycota. Microbiol Spectr. 2017; 5(3):1–29. https://dori.org/10.1128/microbiolspec.funk-0046-2016. PMID: 28597825.

38. Nieuwenhuis BPS, Billiard S, Vuilleumier S, Petit E, Hood ME, Giraud T. Evolution of uni- and bifactorial sexual compatibility systems in fungi. Heredity. 2013; 111:445–55. 10.1038/hdy.2013.67 PMID: 23838688.

39. Kües U, Casselton LA. Homeodomains and regulation of sexual development in basidiomycetes. Trends in Genetics. 1992; 8:154–5. 10.1016/0168-9525(92)90207-k PMID: 1369739.

40. David-Palma M, Sampaio JP, Gonçalves P. Genetic dissection of sexual reproduction in a primary homothallic Basidiomycete. PLoS Genetics. 2016; 12(6):e1006110. 10.1371/journal.pgen.1006110 PMID: 27327578.

41. Hull CM, Boily MJ, Heitman J. Sex-specific homeodomain proteins Sxi1alpha and Sxi2a coordinately regulate sexual development in *Cryptococcus neoformans*. Eukaryot Cell. 2005; 4:526–35. 10.1128/EC.4.3.526-535.2005 PMID: 15755915.

42. Yi R, Mukaiyama H, Tachikawa T, Shimomura N, Aimi T. 2010. A-mating type gene expression can drive clamp cell formation in the bipolar mushroom *Pholiota microspora* (*Pholiota nameko*). Eukaryotic Cell 9: 1109–19. 10.1128/EC.00374-09 PMID: 20453073.

43. Wahl R, Zahiri A, Kämper J. The *Ustilago maydis b* mating type locus controls hyphal proliferation and expression of secreted virulence factors in planta. Molecular Microbiology. 2010; 75(1):208–20. 10.1111/j.1365-2958.2009.06984.x PMID: 19943901.

44. Raudaskoski M, Kothe E. Basidiomycetes mating type gens and pheromone signaling. Eukaryotic Cell. 2010; 9:847–59. 10.1128/EC.00319-09 PMID: 20190072.

45. Duboule D. Guidebook to the Homeobox Genes. Oxford University Press. New York, 1994; pp. 27–71.

46. Liu Y, Steenkamp E T, Brinkmann H, Forget L, Philippe H, Lang BF. Phylogenomic analyses predict sister group relationship of nucleariids and fungi and paraphyly of zygomycetes with significant support. BMC Evolution Biology. 2009; 25:272. 10.1186/1471-2148-9-272 PMID: 19939264.

47. Zuo SX. Study on virulence variation of *Puccinia striiformis* f. sp. *tritici* through sexual hybridization between *P. striiformis* f. sp. *tritici* and *P. striiformis* f. sp. *leymi*. Master Dissertation, Northwest A & F University, Yangling, China. 2018.

48. Zheng D. Sexual hybridization of *Puccinia striiformis* f. sp. *tritici* and *P. striiformis* f. sp. *elymi* in virulence variation of *Puccinia striiformis* f. sp. *tritici*. Master Dissertation, Northwest A & F University, Yangling, China. 2018.

49. Ma XY. Sexual genetic recombination between *Puccinia striiformis* f. sp. *tritici* and *P. achnatheri-sibirici*. Master Dissertation, Northwest A & F University, Yangling, China. 2020.

50. Cheng JJ, Kang ZS, Huang LL, Wang MN, Wan AM. Cheng P. Molecular identification of somatic recombination of stripe rust pathogen (*Puccinia striiformis*) in the greenhouse. Mycosystema. 2015; 34:1128–42. 10.13346/j.mycosystema.140229.

51. Cheng P, Chen XM. Somatic hybridization in *Puccinia striiformis* revealed by virulence patterns and microsatellite markers. Phytopathology. 2009; 99:S23.

52. Holden S, Bakkeren G, Hubensky J, Bamrah R, Abbasi M, Qutob D, de Graaf ML, Kim SH, Kutcher HR, McCallum BD, Randhawa HS, Iqbal M, Uloth K, Burlakoti RR, Brar GS. Uncovering the history of recombination and population structure in western Canadian stripe rust populations through mating type alleles. BMC Biology. 2023; 21:233. 10.1186/s12915-023-01717-9 PMID: 37880702.

53. Kües U. From two to many: multiple mating types in basidiomycetes. Fungal Biology Reviews. 2015; 29:126–66. 10.1016/j.fbr.2015.11.001.

54. Cuomo CA, Bakkeren G, Khalil HB, Panwar V, Joly D, Linning R, Sakthikuma S, Song X, Adiconis X, Fan L, Goldberg JM, Levin JZ, Young S, Zeng QD, Anikster YS, Bruce M, Wang MN, Yin CT, McCallum B, Szabo LJ, Hulbert S, Chen XM, Fellers JP. Comparative analysis highlights variable genome content of wheat rusts and divergence of the mating loci. G3: Genes|Genomes|Genetics. 2017; 7:361–76. 10.1534/g3.116.032797 PMID: 27913634.

55. Zhao J, Zhao SL, Peng YL, Qin JF, Huang LL, Kang ZS. Investigation on geographic distribution and identification of six *Berberis* spp. serving as alternate host for *Puccinia striiformis* f. sp. *tritici* in Linzhi, Tibet. Acta Phytopathologica Sinica. 2016; 46:103–11. 10.13926/j.cnki.apps.2016.01.012.

56. Du ZM, Yao Q, Huang SJ, Yan JH, Hou L, Guo QY, Zhao J, Kang ZS. Investigation and identification of barberry as alternate hosts for *Puccinia striiformis* f. sp. *tritici* in eastern Qinghai. Acta Phytopathologica Sinica. 2019; 49:370–8. 10.13926/j.cnki.apps.000290.

57. Zhuang H, Zhao J, Huang LL, Kang ZS, Zhao J. Identification of three *Berberis* species as potential alternate hosts for *Puccinia striiformis* f. sp. *tritici* in wheat-growing regions of Xinjiang, China. Journal of Integrative Agriculture. 2019; 18:2786–92. 10.1016/S2095-3119(19)62709-7.

58. Li SN, Chen W, Ma XY, Tian XX, Liu Y, Huang LL, Kang ZS, Zhao J. Identification of eight *Berberis* species from the Yunnan-Guizhou plateau as aecial hosts for *Puccinia striiformis* f. sp. *tritici*, the wheat stripe rust pathogen. Journal of Integrative Agriculture. 2021; 20:1563–9. 10.1016/S2095-3119(20)63327-5.

59. Tian Y, Zhan GM, Chen XM, Tungruentragoon A, Lu X, Zhao J, Huang LL, Kang ZS. Virulence and simple sequence repeat marker segregation in a *Puccinia striiformis* f. sp. *tritici* population produced by selfing a Chinese isolate on *Berberis shensiana*. Phytopathology. 2016; 106:185–91. 10.1094/PHYTO-07-15-0162-R PMID: 26551448.

60. Tian Y, Zhan GM, Lu X, Zhao J, Huang LL, Kang ZS. Determination of heterozygosity for avirulence/virulence loci through sexual hybridization of *Puccinia striiformis* f. sp. *tritici*. Frontiers of Agricultural Science and Engineering. 2017; 4:48–58. 10.15302/J-FASE-2016114.

61. Yuan CY, Wang MN, Skinner DZ, See DR, Xia CJ, Guo XH, Chen XM. Inheritance of virulence, construction of a linkage map, and mapping dominant virulence genes in *Puccinia striiformis*f. sp. *tritici* through characterization of a sexual population with genotyping by-sequencing.Phytopathology. 2017; 108:133–41. 10.1094/PHYTO-04-17-0139-R PMID:28876207.

62. Wang L, Zheng D, Zuo SX, Chen XM, Zhuang H, Huang LL, Kang ZS, Zhao J. Inheritance and linkage of virulence genes in Chinese predominant race CYR32 of the wheat stripe rust pathogen *Puccinia striiformis* f. sp. *tritici*. Frontiers in Plant Science. 2018; 9:120. 10.3389/fpls.2018.00120 PMID: 29472940.

63. Kuang WJ, Zhang ZY, Ji HL, Xiang YJ, Zhang M, Peng YL. Population diversity of *Puccinia striiformis* in Linzhi, Tibet. Southwest China Journal of Agricultural Sciences. 2012; 25:1668–73. 10.16213/j.cnki.scjas.2012.05.010.

64. Hu XP, Ma LJ, Liu TG, Wang CH, Peng YL, Pu Q, Xu XM. Population genetic analysis of *Puccinia striiformis* f. sp. *tritici* suggests two distinct populations in Tibet and the other regions of China. Plant Disease. 2017; 101:288–96. 10.1094/PDIS-02-16-0190-RE PMID: 30681929.

65. Wan Q, Liang JM, Luo Y, Ma ZH. Population genetic structure of *Puccinia striiformis* in Northwestern China. Plant Disease. 2015; 99:1764–74. 10.1094/PDIS-02-15-0144-RE PMID: 30699507.

66. Awais M, Ma JB, Chen WB, Zhang BB, Turakulov KS, Li L, Egamberdieva D, Karimjonovich MS, Kang ZS, Zhao J. Molecular genotyping revealed the gene flow of *Puccinia striiformis* f. sp. *tritici* clonal lineage from Uzbekistan of Central Asia to Xinjiang of China. Phytopathology Research. 2025; 7:2. 10.1186/s42483-024-00290-5

66. Jiang B, Wang C, Guo C, Lv X, Gong W, Chang J, He H, Feng J, Chen X, Ma Z. Genetic relationships of *Puccinia striiformis* f. sp. *tritici* in southwestern and northwestern China. Microbiology Spectrum. 2022; 10(4):e0153022. 10.1128/spectrum.01530-22 PMID: 35894618.

67. Hu X, Fu S, Li Y, Xu X, Hu X. Dynamics of *Puccinia striiformis* f. sp. *tritici* urediniospores in Longnan, a critical oversummering region of China. Plant Disease. 2023;107(10):3155–63. 10.1094/PDIS-02-23-0237-RE PMID: 37163309.

68. Li FQ, Han DJ, Wie GR, Zeng QD, Huang LL, Kang ZS. Molecular detection of stripe rust resistant genes in 126 winter wheat varieties from the Huanghuai wheat region. Scientia Agricultura Sinica. 2008; 41:3060–9.

69. Wang H, Yang XB, Ma Z. Long-Distance Spore Transport of Wheat Stripe Rust Pathogen from Sichuan, Yunnan, and Guizhou in Southwestern China. Plant Disease. 2010; 94(7):873–80. 10.1094/PDIS-94-7-0873 PMID: 30743545.

70. Wang FL, Wu LR, Meng QY, Lai SL, Xie SX. Primarily study on new isolates of *Puccinia striiformis* f. sp. *tritici*. Plant Protection. 1992; 18:2–4.

71. Liu TG, Peng YL, Chen WQ, Zhang ZY. First detection of virulence in *Puccinia striiformis* f. sp. *tritici* in China to resistance genes *Yr24* (= *Yr26*) present in wheat cultivar Chuanmai 42. Plant Disease. 2010; 94:1163. 10.1094/PDIS-94-9-1163C PMID: 30743708.

72. Lu NH, Wang JF, Chen XM, Zhan GM, Chen CQ, Huang LL, Kang ZS. Spatial genetic diversity and interregional spread of *P. striiformis* f. sp. *tritici* in Northwest China. European Journal of Plant Pathology. 2011; 131:685–93. 10.1007/s10658-011-9842-y.

73. Hovmøller MS, Justesen AF. Rates of evolution of avirulence phenotypes and DNA markers in a northwest European population of *Puccinia striiformis* f. sp. *tritici*. Molecular Ecology. 2007; 16:4637–47. 10.1111/j.1365-294X.2007.03513.x PMID: 17887968.

74. Liu TL, Bai Q, Wang MN, Li YX, Wan AM, See DR, Xia CJ, Chen XM. Genotyping *Puccinia striiformis* f. sp. *tritici* isolates with SSR and SP-SNP markers reveals dynamics of the wheat stripe rust pathogen in the United States from 1968 to 2009 and identifies avirulence-associated markers. Phytopathology. 2021; 111:1828–39. 10.1094/PHYTO-01-21-0010-R PMID: 33720751.

75. Ding Y, Cuddy WS, Wellings CR, Zhang P, Thach T, Hovmøller MS, Qutob D, Brar GS, Kutcher HR, Park RF. Incursions of divergent genotypes, evolution of virulence and host jumps shape a continental clonal population of the stripe rust pathogen *Puccinia striiformis*. Mol Ecol. 2021; 30(24):6566–84. 10.1111/mec.16182 PMID: 34543497.

76. Wellings CR. *Puccinia striiformis* in Australia: a review of the incursion, evolution, and adaption of stripe rust in the period 1979-2006. Australian Journal of Agricultural Research. 2007; 58:567–75. 10.1071/AR07130.

77. Li Q, Qin JF, Zhao YY, Zhao J, Huang LL, Kang ZS. Virulence analysis of sexual progeny of the wheat stripe rust pathogen recovered from wild barberry in Shaanxi and Gansu. Acta Phytopathologica Sinica. 2016; 46:809–20. 10.13926/j.cnki.apps.2016.06.011.

78. Berlin A, Kyaschenko J, Justesen AF, Jin Y. Rust fungi forming aecia on *Berberis* spp. in Sweden. Plant Disease. 2013; 97:1281–7. 10.1094/PDIS-10-12-0989-RE PMID: 30722146.

79. Wang MN, Chen XM. First report of Oregon grape (*Mahonia aquifolium*) as an alternate host for the wheat stripe rust pathogen (*Puccinia striiformis* f. sp. *tritici*) under artificial inoculation. Plant Disease. 2013; 97:839. 10.1094/PDIS-09-12-0864-PDN PMID: 30722629.

80. Zhang GS, Liu W, Wang L, Ju M, Tian X, Du ZM, Kang Z S, Zhao J. Genetic characteristics and linkage of virulence genes of the *Puccinia striiformis* f. sp. *tritici* isolate TSA-6 to *Yr5* host resistance. Plant Disease. 2023; 107(3):688–700. 10.1094/PDIS-07-22-1637-RE PMID: 35869586.

81. Aljanabi SM, Martinez I. Universal and rapid salt-extraction of high quality genomic DNA for PCR-based techniques. Nucleic Acids Research. 1997; 25:4692–3. 10.1093/nar/25.22.4692 PMID: 9358185.

82. Line R F, Qayoum A. Virulence, aggressiveness, evolution, and distribution of races of Puccinia striiformis (the cause of stripe rust of wheat) in North America, 1968–87. US Department of Agriculture, Agriculture Research Services, Technical Bulletin, 1788. 1992; 44 pp.

83. Pritchard JK, Stephens M, Donnelly P. Inference of population structure using multilocus genotype data. Genetics. 2000; 155:945–59. 10.1093/genetics/155.2.945 PMID: 10835412.

84. Jakobsson M, Rosenberg NA. CLUMPP: a cluster matching and permutation program for dealing with label switching and multimodality in analysis of population structure. Bioinformatics. 2007; 23:1801–6. 10.1093/bioinformatics/btm233 PMID: 17485429.

85. Jombart T. Adegenet: a R package for the multivariate analysis of genetic markers. Bioinformatics. 2008; 24:1403–5. 10.1093/bioinformatics/btn129 PMID: 18397895.

86. Yeh FC, Yang R, Boyle T. POPGENE version 1.32: Microsoft windows-based freeware for population genetic analysis, a quick user guide. University of Alberta, Center for International Forestry Research, Alberta, Canada. 1999.

87. Nei M. Genetic distance between populations. The American Naturalist. 1972; 106:283–92.

